# The Epigenetic Regulator Histone Demethylase KDM5A is Activated and Pathogenic in a Mouse Model of Heart Failure

**DOI:** 10.64898/2026.01.05.697798

**Authors:** Manisha Deogharia, Chitra Lekha Reddy, Ritapa Chaudhuri, Manogna Sai Chirasani, Abhinav K Jain, Leila Rouhi, A J Marian, Priyatansh Gurha

**Affiliations:** Center for Cardiovascular Genetics, Institute of Molecular Medicine, University of Texas Health Sciences Center at Houston, Houston, TX 77030; Department of Epigenetics and Molecular Carcinogenesis, Center for Cancer Epigenetics, The University of Texas MD Anderson Cancer Center, 1515 Holcombe Blvd, Houston, TX 77030, USA

**Author notes:** Address for Correspondence: Priyatansh Gurha, PhD, Center for Cardiovascular Genetics, Brown Foundation Institute of Molecular Medicine The University of Texas Health Science Centre, 1825 Pressler Street, SRB.

## Abstract

Heart failure is a complex disease characterized by the dysregulation of gene expression that culminates in cardiac dysfunction. Epigenetic regulators play a critical role in control of transcription and are increasingly implicated in the pathogenesis of heart failure. We recently showed that the histone demethylase KDM5A is reactivated in the heart of human patients and mouse models of heart failure. However, its pathogenic role in heart failure remained unknown. Utilizing a mouse model of heart failure caused by the deletion of the *Lmna* gene in cardiomyocytes and referred to as LMNA-cardiomyopathy (LMNA-CMP), we show that KDM5A is activated, and expression levels of genes involved in myocyte structure and function, including oxidative phosphorylation (OXPHOS), are suppressed. To determine the pathogenic role of KDM5A in heart failure, the *Kdm5a* gene was specifically deleted in cardiomyocytes in LMNA-CMP mice (*Myh6-Cre:Lmna*^F/F^:*Kdm5a*^F/F^). The deletion of the *Kdm5a* gene improved cardiac function, prolonged survival, attenuated fibrosis, and reduced cell death in LMNA-CMP mice. Transcriptome analysis showed that the deletion of the *Kdm5a* gene restored the expression of over 1,400 dysregulated genes, including those involved in fatty acid metabolism, myogenesis, and OXPHOS in the *Myh6-Cre:Lmna*^F/F^:*Kdm5a*^F/F^ mice. Genome-wide profiling of the epigenetic histone mark H3 lysine 4 trimethylation (H3K4me3), the main target of KDM5A, by CUT&RUN assay showed that deletion of the *Kdm5a* gene partially restored H3K4me3 deposits at loci encoding cardiac transcription factors and metabolic regulators, including *Tbx5* and *Esrrg*, with concomitant rescue of their downstream targets. These findings identify KDM5A as a key epigenetic regulator of cardiomyocyte gene expression and uncover a mechanistic role for KDM5A in the pathogenesis of heart failure.

## Introduction

Heart failure (HF) is a major cause of mortality and morbidity worldwide^1,2^. Regardless of the underlying etiology, HF development and progression share several hallmark features, including mitochondrial dysfunction, metabolic dysregulation, and alterations in sarcomere composition ^3–7^. These pathological changes are often preceded and/or accompanied by extensive molecular remodeling, which affects cardiac myocyte structure and function and are regulated by epigenetic modifications ^8–12^. Establishment of chromatin/epigenetic changes is mediated by networks of histone-modifying enzymes that write, read, and erase histone marks in a locus and cell type–specific manner ^13,14^. Despite advances, key challenges remain in identifying effective epigenetic targets and clarifying how distinct regulators drive HF pathogenesis. Given the heterogeneity of HF, defining core molecular mechanisms involved in its pathogenesis could serve as additional therapeutic targets for interventions.

The histone H3 lysine 4 trimethylation (H3K4me3), a ubiquitous epigenetic mark, is typically enriched at the promoters of active genes and is a major determinant of gene expression. It is dynamically regulated by histone methyltransferases and demethylases^13–15^. The KDM5 family of histone demethylases are evolutionarily conserved members of the JUMONJI family and the primary regulator of the H3K4 methylation state. The KDM5 family of proteins catalyzes the removal of methyl groups from H3K4me3/2 and thereby acts as a repressor of gene expression, though evidence also supports roles in transcriptional activation through recruitment of co-regulators^13,16–18^.

We recently showed increased KDM5A expression and activity in human HF samples and in mouse models of HF^19,20^. However, the functional and biological significance of activation of the KDM5A in cardiomyocytes in heart failure remains unknown. Therefore, to determine the pathogenic role of KDM5A in HF, we utilized a well-characterized mouse model of HF generated upon the deletion of the *Lmna* gene specifically in cardiac myocytes (*Myh6*-*Cre*:*Lmna*^F/F,^ aka LMNA-CMP mice)^9,21^. These mice exhibit premature death within 3 to 4 weeks after birth, severe cardiac dysfunction, and increased myocardial fibrosis and apoptosis.

Our results show that KDM5A protein levels are increased in cardiac myocytes in the LMNA-CMP and its target genes involved in cardiac structure and function, and OXPHOS are suppressed. Genetic deletion of the *Kdm5a* gene specifically in cardiac myocytes restores expression of the genes and transcription factors involved in cardiac structure and function, and mitochondrial OXPHOS in the LMNA-CMP mice. The findings were partially corroborated by increased genome-wide H3K4me3 methylation of the corresponding genes and transcription factor loci. Partial restoration of the molecular remodeling of the myocardium was associated with prolongation of survival, improved cardiac function, and reduced myocardial fibrosis and cell death. Thus, our study uncovered a key role for KDM5A-mediated epigenetic regulation in mitochondrial homeostasis and cardiac function in heart failure.

## Results

### KDM5A is induced, and its targets are suppressed in cardiomyocytes in LMNA-CMP

The expression of the KDM5 family of proteins, including KDM5A, progressively declines in the heart in the post-natal period ^22^. In contrast, in a mouse model of heart failure generated upon deletion of the *Lmna* gene (*Lmna*^−/−^), KDM5A is markedly induced in the postnatal heart, and the expression of its target genes is suppressed ^20^. To investigate the role of KDM5A in the pathogenesis of heart failure, we used a cardiomyocyte-specific LMNA cardiomyopathy mouse model in which the *Lmna* gene is selectively deleted in cardiomyocytes using the *Myh6-Cre* system (*Myh6-Cre:Lmna*^F/F^) (9). These mice develop cardiac dysfunction characterized by marked cardiomyocyte apoptosis and uniform mortality by approximately four weeks of age (9). Heterozygous deletion of the *Lmna* gene results in a similar cardiac phenotype but evolves more gradually, with disease progression occurring over a 6–9-month period^9,23,24^. Given the shared phenotypic features but an accelerated course, the homozygous cardiomyocyte-specific *Lmna* knockout model was selected for the studies.

Analysis of whole-heart lysates from *Myh6-Cre:Lmna*^F/F^ mice by immunoblotting (IB) showed increased KDM5A expression level compared to WT controls (Figures 1A and 1B). Similarly, the KDM5A protein level was increased in cardiomyocytes isolated from the *Myh6-Cre:Lmna*^F/F^ mice as compared to the WT myocytes (Figures 1C and 1D). Given that KDM5A is a histone demethylase predominantly localized to the nucleus, the expression level of KDM5A in the cardiomyocyte nuclear protein extracts was analyzed. IB analysis showed an increased level of KDM5A in *Myh6-Cre:Lmna*^F/F^ mice (Figures 1E and 1F). Likewise, transcript levels of KDM5A putative target genes involved in oxidative phosphorylation (OXPHOS), fatty acid oxidation (FAO), including ERR family of transcription factors, were reduced in the *Myh6-Cre:Lmna*^F/F^ cardiomyocytes (Figure 1G). Taken together, these findings demonstrate that KDM5A levels, including in the myocyte nuclei, are increased and its downstream targets, including those involved in metabolism and myocyte structure, are suppressed in a mouse model of heart failure.

**Figure 1:**
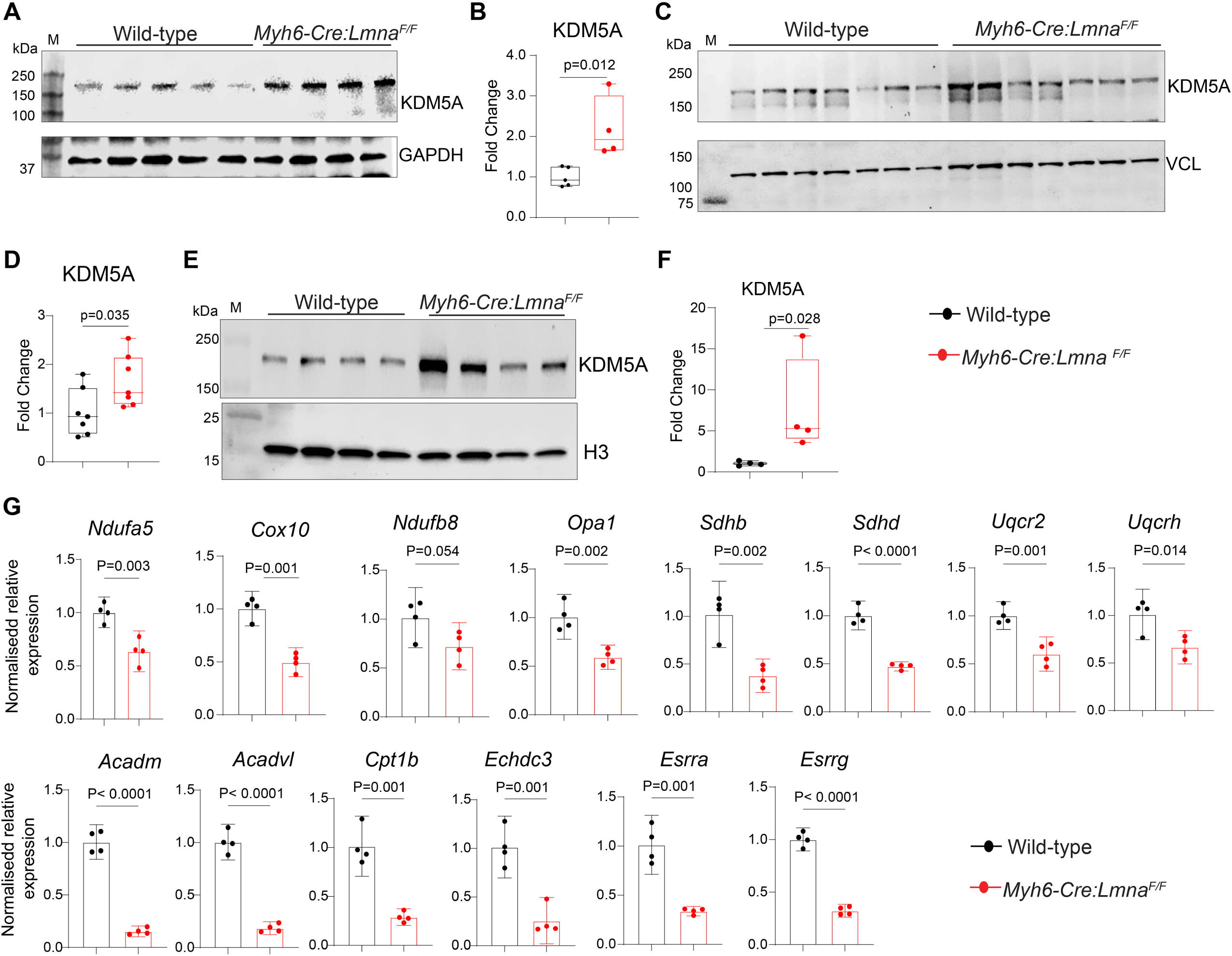
Expression of KDM5A protein is increased and its target genes are suppressed in *Myh6-Cre:Lmna*^F/F^ cardiomyocytes. **(A)** Immunoblot analysis of whole-heart lysates from Wild-type and *Myh6-Cre:Lmna*^F/F^ mice showing KDM5A protein expression and normalization control GAPDH. **(B)** Quantification of KDM5A protein levels normalized to GAPDH, demonstrating increased expression in *Myh6-Cre:Lmna*^F/F^ hearts compared to Wild-type (n=5 Wild-type and n=4 *Myh6-Cre:Lmna*^F/F^). **(C)** Immunoblot analysis of isolated cardiomyocytes from Wild-type and *Myh6-Cre:Lmna*^F/F^ mice showing KDM5A expression. **(D)** Quantification of KDM5A protein levels from isolated cardiomyocytes normalized to Vinculin (VCL), confirming increased expression in *Myh6-Cre:Lmna*^F/F^ samples (n=7 Wild-type and *Myh6-Cre:Lmna*^F/F^). **(E)** Immunoblot analysis of nuclear extracts from isolated cardiomyocytes of Wild-type and *Myh6-Cre:Lmna*^F/F^ mice. **(F)** Quantification of nuclear KDM5A protein levels normalized to Histone H3, showing enhanced nuclear retention in *Myh6-Cre:Lmna*^F/F^ cardiomyocytes (n=4 Wild-type and *Myh6-Cre:Lmna*^F/F^). **(G)** Transcript levels of representative KDM5A target genes in cardiomyocytes, showing suppression of genes in *Myh6-Cre:Lmna*^F/F^ samples as compared to Wild-type (n=4 Wild-type and *Myh6-Cre:Lmna*^F/F^). All data are presented as mean ± 95% confidence interval (CI). Normality of data distribution was assessed using the Shapiro–Wilk test. Statistical significance was evaluated using the Student’s *t*-test for normally distributed variables (Panel B, D and G) or the Mann–Whitney *U* test for non-normally distributed variables (Panel F). The corresponding *p*-values are shown.

### *Kdm5a* deletion improves survival and cardiac function in the LMNA-CMP mice

To elucidate the role of KDM5A induction in heart failure, we generated cardiomyocyte-specific *Kdm5a* single knockout mice (*Myh6-Cre:Kdm5a*^F/F^). Both heterozygous and homozygous *Kdm5a* knockout offspring were born at the expected Mendelian ratios and exhibited no discernible phenotypes at birth. The efficiency of *Kdm5a* deletion was assessed by immunoblot analysis of cardiomyocyte lysates from WT, *Kdm5a^F/F^,* and *Myh6-Cre:Kdm5a^F/F^* mice. KDM5A protein levels were significantly reduced in the knockout mouse hearts (Figure 2 A-B).

**Figure 2.**
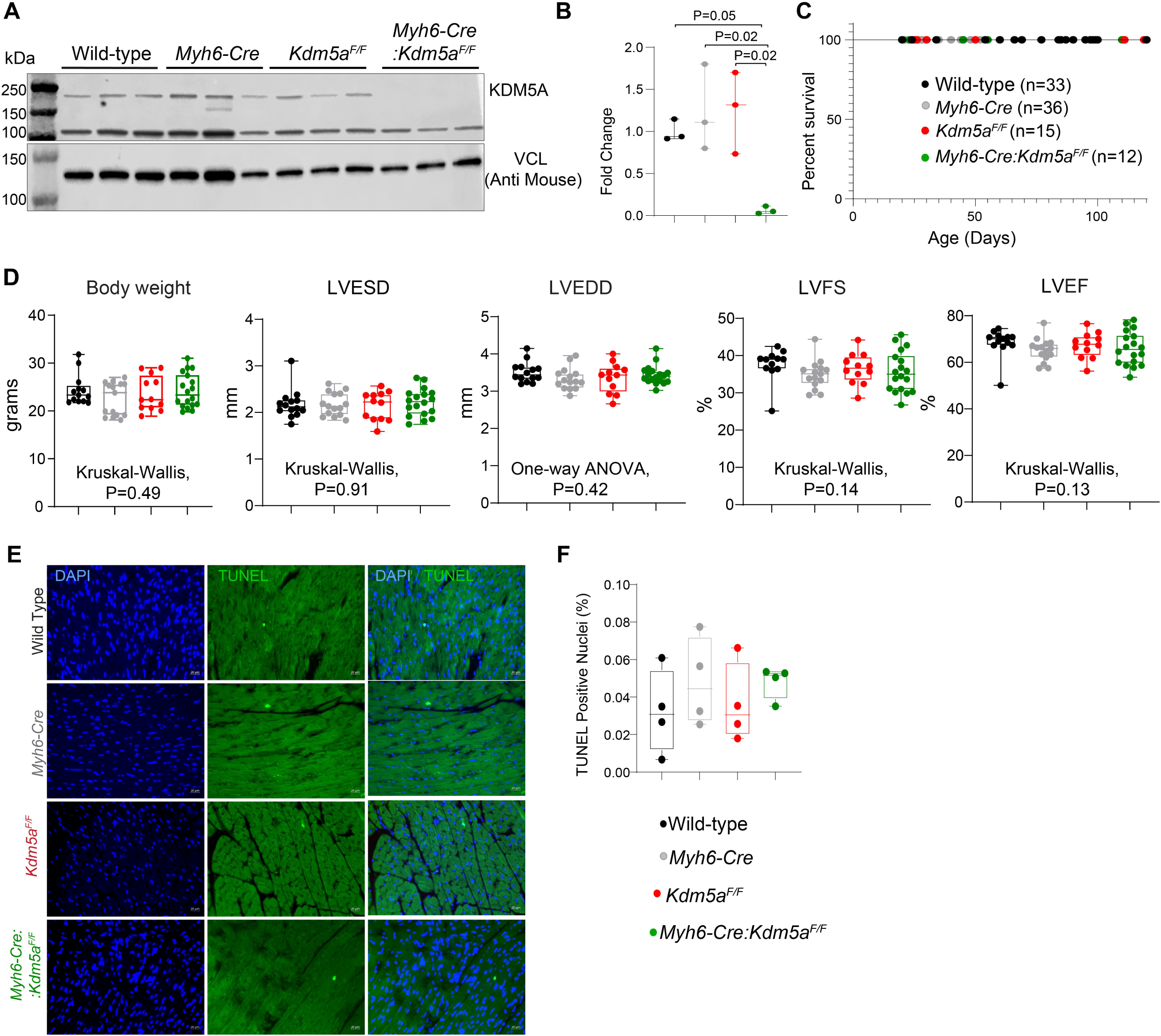
Conditional deletion of *Kdm5a* in cardiomyocytes does not cause overt phenotypic abnormalities. (**A-B)** Immunoblot analysis of isolated cardiomyocytes from P16 mice, which confirmed the absence of KDM5A protein following cardiomyocyte-specific deletion, normalized to Vinculin (VCL) as a loading control (n=3). **(B)** Quantitative assessment of KDM5A deletion. Statistical analysis was performed by Kruskal-Wallis test and Dunn post hoc pairwise comparisons P values are shown. **(C)** Kaplan–Meier survival analysis showing that loss of *Kdm5a* in cardiomyocytes has no significant effect on overall survival compared with control groups. **(D)** Echocardiographic measurements obtained from four groups of mice assessing body weight, heart rate, left ventricular end-diastolic diameter (LVEDD), left ventricular end-systolic diameter (LVESD), and fractional shortening (FS). No significant differences were observed across genotypes. Data are presented as mean ± 95% confidence interval (CI). Statistical significance was evaluated by one-way ANOVA followed by Tukey’s post hoc pairwise comparisons or Kruskal-Wallis test and Dunn post hoc pairwise comparison. The one-way ANOVA or Kruskal-Wallis test P values are shown in the figure. **(E)** TUNEL staining of myocardial sections from 2-month-old mice (n = 4 per group) showing apoptotic nuclei (green) and DAPI (blue). **(F)** Quantification of TUNEL^+^ nuclei expressed as percentage of total nuclei, showing no difference between the groups (n=4,mean ± SD, one-way ANOVA with post hoc Tukey pairwise comparison).

To determine the effect of *Kdm5a* deletion on survival, Kaplan–Meier survival analyses were performed, and cardiac function was assessed by transthoracic echocardiography. *Myh6-Cre* mice were included as controls to account for potential nonspecific effects of Cre recombinase and exhibited no detectable cardiac phenotype when evaluated at 2 months of age. Cardiomyocyte-specific deletion of *Kdm5a* had no significant effect on cardiac function or survival up to 120 days of age, as shown in Figure 2C and 2D and summarized in Table 1. Likewise, no increase in apoptosis was observed in any group, as assessed by TUNEL staining and quantitative analysis (Figure 2E and F). Our findings suggest that *Kdm5a* suppression is well tolerated in adult mice, consistent with its developmental downregulation in the post-natal period. These findings enable the determination of the effects of KDM5A activation on cardiac pathology in LMNA-CMP.

To evaluate the impact of *Kdm5a* deletion in LMNA cardiomyopathy, we generated cardiomyocyte-specific double-knockout mice (*Myh6-Cre:Lmna*^F/F^:*Kdm5a*^F/F^). Because cardiomyocyte-specific deletion of *Kdm5a* alone did not cause a measurable abnormality in cardiac structure, function, survival, or apoptosis, *Kdm5a* knockout-only group was included in downstream analyses.

*Kdm5a* deletion did not affect body weight in the *Myh6-Cre:Lmna^F/F^*mice. Deletion of the *Lmna* gene, however, was associated with reduced body weight at 3 weeks of age, regardless of the *Kdm5a* deletion (Figure 3A and Table 2). We next assessed cardiac size and function by echocardiography at 3 weeks of age in the WT, *Myh6-Cre:Lmna^F/F^*, and *Myh6-Cre:Lmna^F/F^:Kdm5a^F/F^* mice. As shown previously, the *Myh6-Cre:Lmna^F/F^* mice had severe cardiac dilation and dysfunction^9^. Deletion of the *Kdm5a* gene in the LMNA-CMP mice (*Myh6-Cre:Lmna^F/F^:Kdm5a^F/F^)* significantly improved left ventricular size and function, as evidenced by decreased end-systolic diameters (LVESD), LV end-diastolic (LVEDD), and LVEDD indexes to body weight (LVEDDi), and increased LV fractional shortening (LVFS) and ejection fraction (LVEF, Table 2 and Figure 3D). Kaplan-Meier survival plot showed that deletion of the *Kdm5a* gene extended the median lifespan of *Myh6-Cre:Lmna^F/F^* mice and improved overall survival (χ² =99.1, p<0.0001; Figure 3E). In agreement with these findings, as shown in Figure 3F, deletion of the *Kdm5a* gene attenuated the expression of the molecular markers of cardiac hypertrophy and heart failure in *Myh6-Cre:Lmna^F/F^:Kdm5a^F/F^* CM as determined by quantitative RT-PCR (qRT-PCR).

**Figure 3.**
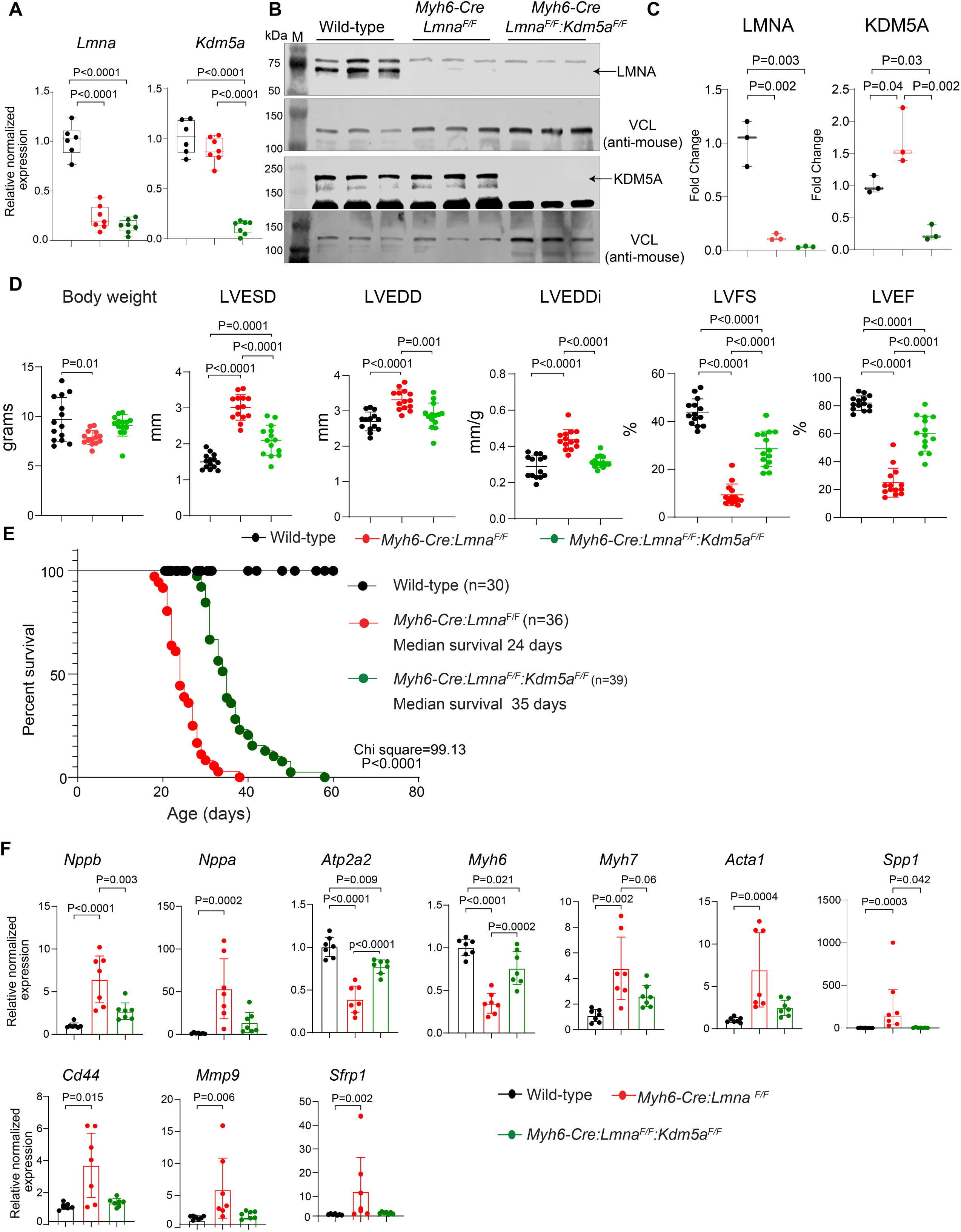
Conditional deletion of *Kdm5a* in mice attenuates cardiac dysfunction and improves survival of LMNA-CMP mice. **(A)** Quantitative RT-PCR showing deletion of mRNA levels of *Lmna* and *Kdm5a* in the cardiac myocytes showing substantial reduction of *Lmna* in *Myh6-Cre:Lmna^F/F^*and *Myh6-Cre:Lmna^F/F^:Kdm5a*^F/F^ and *Kdm5a* in the latter group compared to WT controls. **(B-C)** Immunoblot and quantitative analysis showing reduction of LMNA in the *Myh6-Cre:Lmna^F/F^* and *Myh6-Cre:Lmna^F/F^:Kdm5a*^F/F^ and KDM5A in the *Myh6-Cre:Lmna^F/F^:Kdm5a*^F/F^ group. The data confirmed deletion of both genes. **(D)** Body weight, heart rate, and selected echocardiographic indices were measured in Wild-type, *Myh6-Cre:Lmna^F/F^* and *Myh6-Cre:Lmna^F/F^:Kdm5a*^F/F^ mice (n = 14 per group). Parameters include Body weight, LVESD (left ventricular end-systolic diameter), LVEDD (left ventricular end-diastolic diameter), LVEDDi (LVEDD indexed to body weight), FS (fractional shortening) and EF (ejection fraction). Statistical significance was determined by one-way ANOVA followed by Tukey post hoc test. **(E)** Kaplan–Meier survival curves demonstrating prolonged median and maximal survival in *Myh6-Cre:Lmna^F/F^:Kdm5a*^F/F^ mice compared to *Myh6-Cre:Lmna^F/F^*. **(F)** Transcript levels of selected markers of cardiac dysfunction measured by quantitative RT-qPCR in isolated cardiomyocytes from P21 mice. Expression values were normalized to *Gapdh*. Data are presented as mean ± 95% confidence interval (CI). N= 6-7, Statistical significance was determined using one-way ANOVA followed by Tukey post hoc pairwise comparisons.

### *Kdm5a* deletion attenuates apoptosis in LMNA-CMP

The observed changes in cardiac function led us to analyse the apoptotic pathways, as cell death is an integral phenotype of LMNA-CMP^9,20,25^ and heart failure in general. GSEA showed enrichment of genes involved in apoptosis in *Myh6-Cre:Lmna^F/F^*, as compared to WT cardiac myocytes (Figure 4A). Deletion of the *Kdm5a* gene significantly suppressed pro-apoptotic gene expression signatures in *Myh6-Cre:Lmna^F/F^*:*Kdm5a*^F/F^ mice, as demonstrated by GSEA plots derived from RNA-Seq data (Figure 4A).

**Figure 4:**
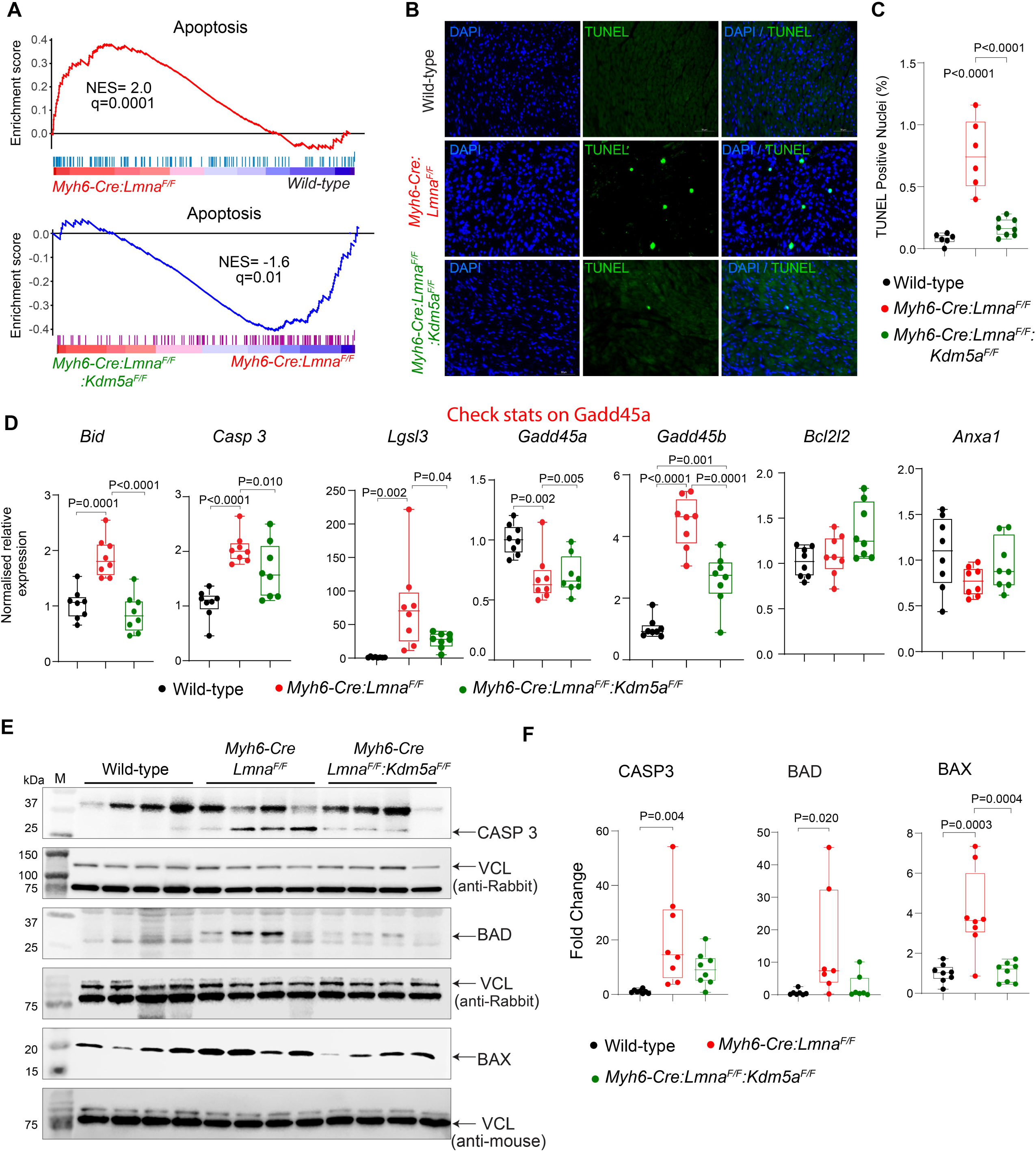
*Kdm5a* deletion attenuates apoptosis in LMNA-CMP hearts. **(A)** Gene set enrichment analysis (GSEA) of RNA-seq data showing significant induction of apoptotic gene signatures in *Myh6-Cre:Lmna*^F/F^ cardiomyocytes and reversal of this signature upon *Kdm5a* deletion in *Myh6-Cre:Lmna*^F/F^:*Kdm5a*^F/F^ CM. **(B)** Representative TUNEL staining of myocardial sections from P21 mice demonstrating increased apoptotic nuclei (green) in *Myh6-Cre:Lmna*^F/F^ hearts, counterstained with DAPI (blue). **(C)** Quantification of TUNEL^+^ nuclei expressed as percentage of total nuclei, showing a marked increase in *Myh6-Cre:Lmna*^F/F^ hearts compared with Wild-type and a significant reduction upon *Kdm5a* deletion. N = 6 per group. Statistical significance was determined using one-way ANOVA followed by Tukey post hoc pairwise comparisons. **(D)** Transcript levels of pro-apoptotic genes measured by quantitative RT-qPCR from whole heart RNA at P21, normalized to *Gapdh*, showing upregulation in *Myh6-Cre:Lmna*^F/F^ and partial restoration in *Myh6-Cre:Lmna*^F/F^:*Kdm5a*^F/F^. N = 7 biological replicates per group were used and Statistical significance was determined using one-way ANOVA followed by Tukey post hoc pairwise comparisons. **(E)** Representative immunoblot (IB) analysis of cardiomyocyte extracts from P21 mice showing expression of CASP3, BAX, and BAD across the three groups. **(F)** Quantification of IB data (normalized to VCL) reveals increased expression of pro-apoptotic proteins in *Myh6-Cre:Lmna*^F/F^ hearts, which are attenuated following *Kdm5a* deletion. N = 8 biological replicates per group were used and Statistical significance was determined using one-way ANOVA followed by Tukey post hoc pairwise comparisons.

To assess the extent of apoptosis, we performed a TUNEL assay, which showed an increased number of TUNEL-positive nuclei in the myocardium of *Myh6-Cre:Lmna^F/F^* mice compared to WT (Figure 4B-C). Conversely, the number of TUNEL-positive nuclei was markedly reduced in *Myh6-Cre:Lmna^F/F^*:*Kdm5a*^F/F^ mice, as shown in Figure 4B and 4C. Furthermore, qRT-PCR analysis of the transcripts corroborated the findings in the experimental groups (Figure 4D). Furthermore, immunoblot analysis of cardiac myocyte protein extracts showed upregulation of Caspase-3 (CASP3), Bcl2-associated agonist of cell death (BAD), and Bcl-2-associated X protein (BAX) proteins in *Myh6-Cre:Lmna^F/F^*, and attenuation of their levels upon deletion of the *Kdm5a* gene (*Myh6-Cre:Lmna^F/F^*:*Kdm5a*^F/F^), as shown in Figure 4E-F. Collectively, these results indicate that expression of KDM5A mediates activation of apoptosis in *Myh6-Cre:Lmna^F/F^*and its genetic ablation mitigates the phenotype.

### *Kdm5a* deletion rescues myocardial fibrosis in *Myh6-Cre:Lmna^F/F^* mice

To determine the effect of *Kdm5a* deletion on cardiac phenotypes, cardiac fibrosis was assessed by staining thin myocardial sections with picrosirius red, and the collagen volume fraction (CVF) was calculated, which is used as a quantitative marker of fibrosis. The CVF was significantly increased in *Myh6-Cre:Lmna^F/F^* hearts, comprising 9.8±3.8% of the myocardium (Figure 5A-B), compared to approximately 1% observed in WT hearts. This increase was notably attenuated in *Myh6-Cre:Lmna^F/F^:Kdm5a^F/F^* mice, as indicated by a significant reduction in CVF to 1.5±0.85% (Figure 5B). Quantitative RT-PCR analyses of transcripts associated with fibrosis revealed increased expression levels of *Tgfb1, Tgfb2,* and *Col1a1* among others in the *Myh6-Cre:Lmna^F/F^*hearts, which were significantly attenuated upon *Kdm5a* deletion in *Myh6-Cre:Lmna^F/F^:Kdm5a^F/F^* hearts (Figure 5C).

**Figure 5.**
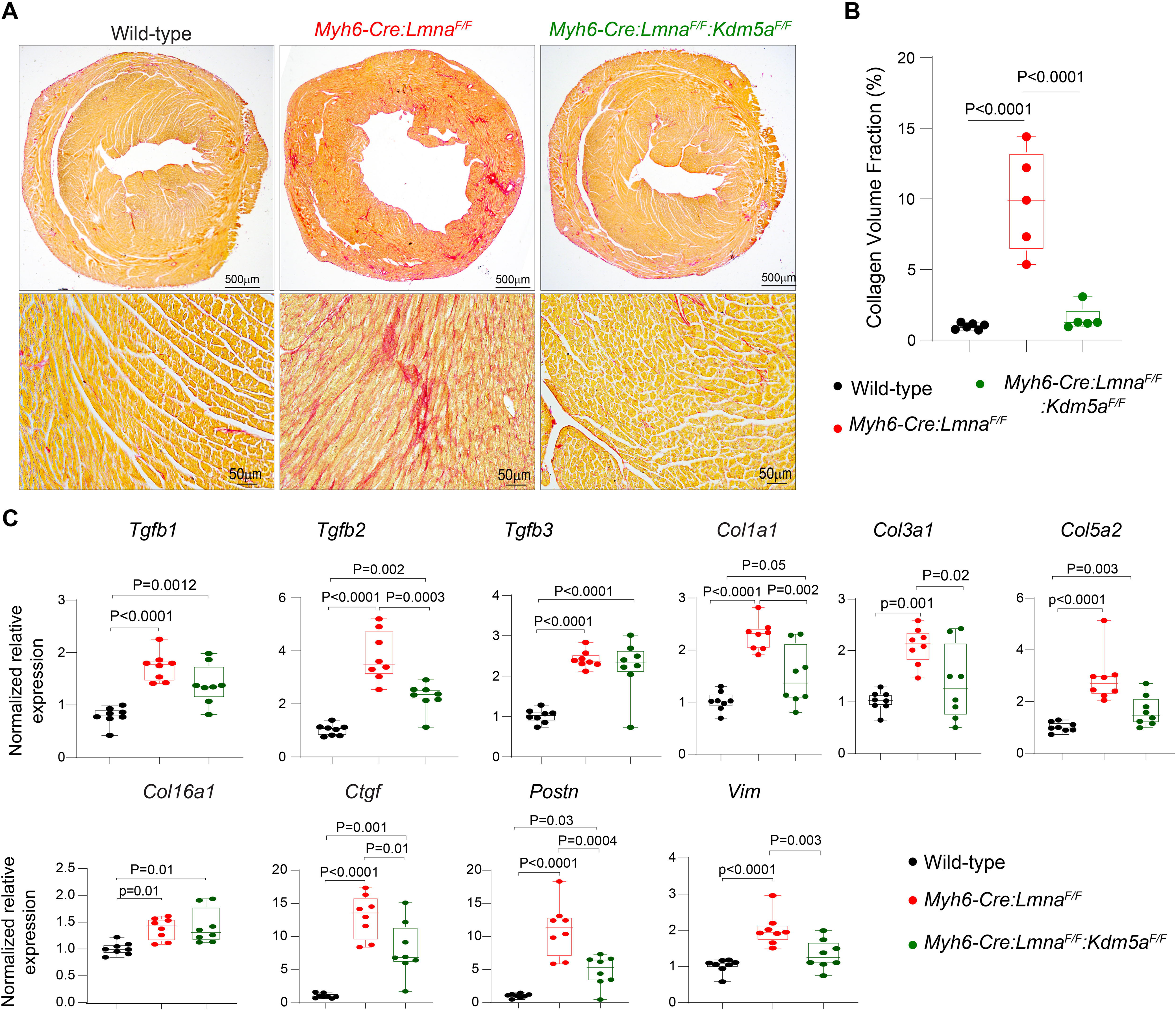
*Kdm5a* deletion rescues myocardial fibrosis in *Myh6-Cre:Lmna*^F/F^ mice. **(A–B)** Representative picrosirius red–stained myocardial sections from 3-week-old Wild-type, *Myh6-Cre:Lmna*^F/F^, and *Myh6-Cre:Lmna*^F/F^:*Kdm5a*^F/F^ mice, with corresponding quantitative analysis of collagen volume fraction (CVF). CVF was markedly elevated in *Myh6-Cre:Lmna*^F/F^ hearts (9.8 ± 3.8%) compared to Wild-type (~1%), and this increase was significantly attenuated in *Myh6-Cre:Lmna*^F/F^:*Kdm5a*^F/F^ hearts (1.5 ± 0.85%). Collagen volume fraction was quantified using ImageJ. For each genotype, n=5 mice hearts were used and statistical significance was determined using one-way ANOVA with Tukey’s post hoc test. **(C)** Transcript levels of fibrosis-associated genes in Wild-type, *Myh6-Cre:Lmna*^F/F^, and *Myh6-Cre:Lmna*^F/F^:*Kdm5a*^F/F^ hearts, measured by quantitative RT-qPCR. N=7 biological replicates per group were used and Statistical significance was determined using one-way ANOVA followed by Tukey post hoc pairwise comparisons.

### Deletion of *Kdm5a* attenuates dysregulation of gene expression in LMNA-CMP

To determine the molecular mechanisms underlying the rescue of heart failure phenotypes upon deletion of the *Kdm5a* gene, we performed RNA-seq analysis on isolated cardiac myocytes from Wild-type, *Myh6-Cre:Lmna^F/F^*, and *Myh6-Cre:Lmna^F/F^:Kdm5a^F/F^* CMs. Principal component analysis (PCA) of the RNA-seq data revealed distinct genotype-dependent segregation of the samples (Figure 6A). Analysis of the RNA-seq data between the *Myh6-Cre:Lmna^F/F^* and WT cardiac myocytes led to the identification of 6,609 differentially expressed genes (DEGs), comprised of 3258 upregulated and 3351 downregulated genes (Figure 6B). Deletion of *Kdm5a* attenuated dysregulation of gene expression in *Myh6-Cre:Lmna^F/F^:Kdm5a^F/F^*cardiac myocytes, as 1,905 genes were differentially expressed compared with *Myh6-Cre:Lmna*^F/F^ (Figure 6C). Of these, 1,458 differentially expressed genes were considered rescued as their expression level was normalized upon *Kdm5a* deletion, i.e., was similar to that in the WT myocytes (Figure 6D). Gene set enrichment analysis (GSEA) of known KDM5A target genes indicated their suppression in the *Myh6-Cre:Lmna^F/F^* and their restoration following the deletion of the *Kdm5a* gene (Figure 6E), which supports the role of KDM5A-mediated epigenetic remodelling in the pathogenesis of heart failure.

**Figure 6:**
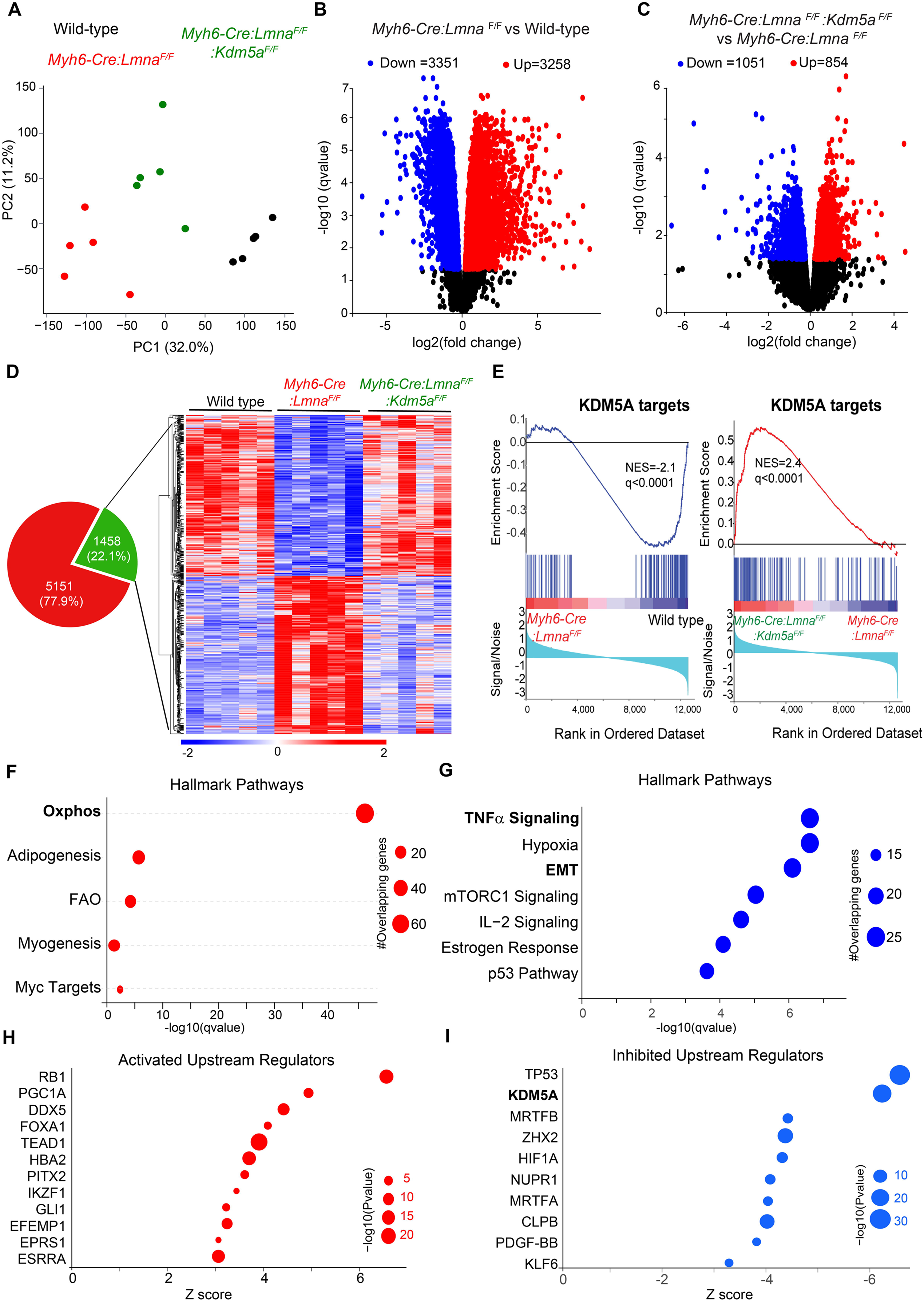
*Kdm5a* deletion in LMNA-CMP restores OXPHOS, FAO, and myogenesis gene expression. **(A)** Principal component analysis (PCA) of RNA-seq data from Wild-type (black), *Myh6-Cre:Lmna*^F/F^ (red), and *Myh6-Cre:Lmna*^F/F^:*Kdm5a*^F/F^ (green) CM showing distinct clustering of transcriptional profiles. Each dot represents an individual biological replicate. **(B)** Volcano plot of differentially expressed genes (DEGs) in *Myh6-Cre:Lmna*^F/F^ vs. Wild-type hearts, highlighting significantly upregulated (red) and downregulated (blue) genes.**(C)** Volcano plot of DEGs in *Myh6-Cre:Lmna*^F/F^:*Kdm5a*^F/F^ vs. *Myh6-Cre:Lmna*^F/F^ CM, showing transcriptional rescue upon *Kdm5a* deletion. **(D)** The Venn diagram depicting the overlap of DEGs showing rescue after *Kdm5a* loss and heatmap of rescue gene showing widespread transcriptional alterations in *Myh6-Cre:Lmna*^F/F^ compared to Wild-type, with partial restoration of expression patterns following *Kdm5a* deletion. **(E)** Gene Set Enrichment Analysis (GSEA) plots for KDM5A target genes. Left: *Myh6-Cre:Lmna*^F/F^ vs. Wild-type, showing significant enrichment of KDM5A targets. Right: *Myh6-Cre:Lmna*^F/F^:*Kdm5a*^F/F^ vs. *Myh6-Cre:Lmna*^F/F^, showing reversal of KDM5A-driven transcriptional repression. **(F)** Hallmark pathway enrichment analysis highlighting rescue of oxidative phosphorylation (OXPHOS), fatty acid oxidation (FAO), and myogenesis pathways upon *Kdm5a* deletion. **(G)** Downregulated hallmark pathways in *Myh6-Cre:Lmna*^F/F^:*Kdm5a*^F/F^ relative to *Myh6-Cre:Lmna*^F/F^, including TNFα signalling, hypoxia, EMT, and p53 pathways. **(H)** IPA predicted upstream regulators that show inhibition in *Myh6-Cre:Lmna*^F/F^:*Kdm5a*^F/F^ compared to *Myh6-Cre:Lmna*^F/F^, including KDM5A, TP53, and HIF1A. **(I)** IPA predicted activated upstream regulators in *Myh6-Cre:Lmna*^F/F^:*Kdm5a*^F/F^ compared to *Myh6-Cre:Lmna*^F/F^, including ESRRA, and PGC1A.

Pathway analysis of genes whose transcripts were rescued upon deletion of the *Kdm5a* gene in LMNA-CMP mice showed a marked upregulation of genes involved in oxidative phosphorylation (OXPHOS) and myogenesis (Figure 6F), whereas those involved in inflammation and fibrosis were suppressed (Figure 6G). KDM5A was the top upstream regulator of the genes whose transcripts were rescued upon deletion of the *Kdm5a* gene. In addition, TP53 and members of the SMAD family were among the upstream regulators whose targets were suppressed upon deletion of the *Kdm5a* gene (Figure 6H); in contrast, targets of Peroxisome proliferator-activated receptor gamma coactivator 1-alpha (PGC1A, Z score=5.1) and the Estrogen-related receptor (ERR, Z score=3.2) were induced (Figure 6I). Collectively, these data demonstrate that KDM5A deletion abrogates the pathogenic transcriptional program in *Myh6-Cre:Lmna^F/F^*myocytes by restoring expression of genes involved in OXPHOS and myogenesis and inducing expression of the key transcription factors, such as the ERR family.

### Genome-wide redistribution of H3K4me3 marks in LMNA-CMP cardiomyocytes and their restoration upon *Kdm5a* deletion

KDM5A preferentially targets cis-regulatory genomic regions that are enriched with H3K4me3 marks for their removal. Therefore, to elucidate the epigenetic changes associated with the deletion of the *Kdm5a* gene, we determined genome-wide H3K4me3 occupancy in cardiac myocytes isolated from WT, *Myh6-Cre:Lmna^F/F,^* and *Myh6-Cre:Lmna^F/F^:Kdm5a^F/F^*mice by CUT&RUN assays. Approximately 15 million uniquely mapped reads per sample were obtained and analyzed, which led to the identification of 21,324 ± 2550 H3K4me3 peaks in WT, 20,042 ± 1917 peaks in *Myh6-Cre:Lmna^F/F^*, and 23,775 ± 2150 peaks in *Myh6-Cre:Lmna^F/F^:Kdm5a^F/F^* samples (Figure 7A). Genomic annotation revealed that differential H3K4me3 peaks were predominantly localized to promoters and transcription start site (TSS)–proximal regions, with additional enrichment at intronic and exonic regulatory elements (Fig. 7B). Overall genomic distribution and peak length of H3K4me3 were comparable across all three genotypes (Fig. 7C). Consistent with canonical H3K4me3 localization, average count per million (CPM) signal profiles demonstrated strong enrichment at TSSs, and heatmap analysis showed robust H3K4me3 signal accumulation within ±3 kb of TSSs in all groups (Fig. 7D and Supplementary Fig. 1). PCA clearly separated WT, *Myh6-Cre:Lmna*^F/F^, and *Myh6-Cre:Lmna*^F/F^:*Kdm5a*^F/F^ samples, indicating genotype-dependent global differences in H3K4me3 landscapes (Fig. 7E).

**Figure 7:**
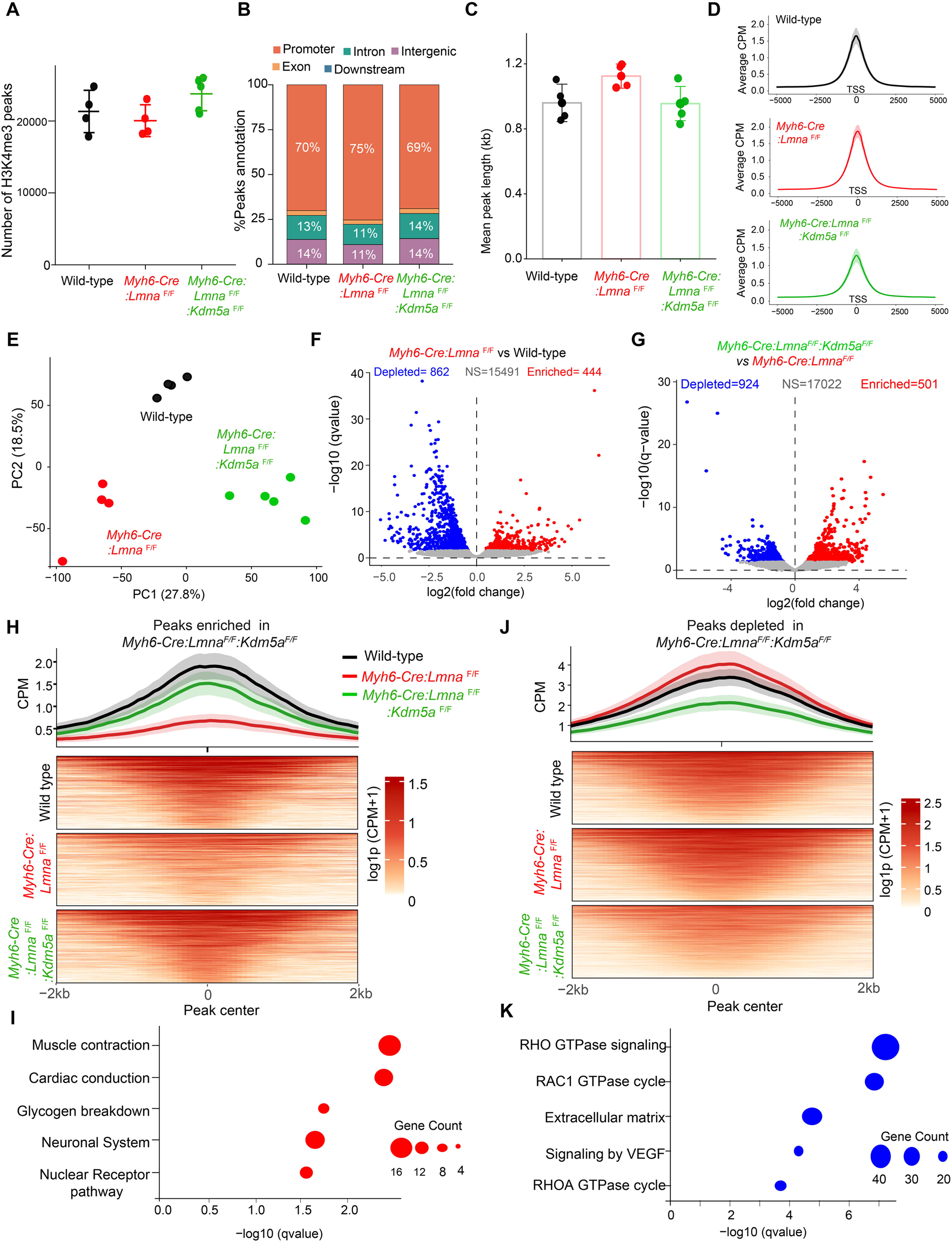
Genome-wide H3K4me3 profiling reveals altered promoter enrichment in LMNA-CMP and rescue upon *Kdm5a* deletion. **(A)** Total number of H3K4me3 peaks identified by CUT&RUN in Wild-type (black), *Myh6-Cre:Lmna*^F/F^ (red), and *Myh6-Cre:Lmna*^F/F^:*Kdm5a*^F/F^ (green) CM. Each dot represents an individual biological replicate; bars indicate mean ± SD. **(B)** Genomic annotation of H3K4me3 peaks across genotypes. **(C)** Mean H3K4me3 peak length per sample, showing similar distribution across genotypes. **(D)** Aggregate H3K4me3 signal profiles centred on genome wide transcription start sites (TSS ±5 kb), demonstrating promoter-proximal H3K4me3 enrichment. **(E)** Principal component analysis (PCA) of genome-wide H3K4me3 profiles separates WT, *Myh6-Cre:Lmna*^F/F^, and *Myh6-Cre:Lmna*^F/F^:*Kdm5a*^F/F^ samples, indicating distinct chromatin states and partial normalization following *Kdm5a* loss. **(F)** Volcano plot showing differentially enriched H3K4me3 peaks in *Myh6-Cre:Lmna*^F/F^ relative to Wild-type. Red, enriched; blue, depleted; grey, not significant. **(G)** Volcano plot of H3K4me3 changes in *Myh6-Cre:Lmna*^F/F^:*Kdm5a*^F/F^ relative to *Myh6-Cre:Lmna*^F/F^, showing restoration of promoter-associated H3K4me3 peaks. **(H)** Average profiles and heatmaps of H3K4me3 signal at peaks enriched in *Myh6-Cre:Lmna*^F/F^:*Kdm5a*^F/F^ relative to *Myh6-Cre:Lmna*^F/F^, demonstrating recovery of promoter H3K4me3 occupancy toward Wild-type levels. **(I)** Gene ontology and pathway enrichment analysis of genes associated with rescued H3K4me3 peaks, revealing restoration of muscle contraction, cardiac conduction, metabolic, and nuclear receptor signalling programs. **(J)** Average profiles and heatmaps of H3K4me3 signal at peaks depleted in *Myh6-Cre:Lmna*^F/F^:*Kdm5a*^F/F^ relative to *Myh6-Cre:Lmna*^F/F^, indicating attenuation of aberrantly elevated promoter marks. **(K)** Pathway enrichment analysis of genes associated with depleted peaks, highlighting downregulation of RHO GTPase signalling, extracellular matrix organization, and VEGF-associated pathways.

Differential peak analysis between *Myh6-Cre:Lmna*^F/F^ and WT cardiomyocytes identified 444 enriched and 862 depleted H3K4me3 peaks in the *Myh6-Cre:Lmna*^F/F^ group (Fig. 7F). Notably, deletion of *Kdm5a* in the *Myh6-Cre:Lmna*^F/F^ background resulted in redistribution of H3K4me3 at 1,425 genomic loci, comprising 501 enriched and 924 depleted peaks in *Myh6-Cre:Lmna*^F/F^:*Kdm5a*^F/F^ cardiomyocytes (Fig. 7G).

Heatmap and meta profile analyses revealed a subset of H3K4me3 peaks that were depleted in *Myh6-Cre:Lmna*^F/F^ cardiomyocytes but restored upon *Kdm5a* deletion (Fig. 7H), suggesting these loci represent direct or primary targets of KDM5A-mediated demethylation. Pathway analysis showed that these rescued H3K4me3 peaks are enriched at genes essential for cardiac function (Figure 7I). Conversely, H3K4me3 peaks that were increased in *Myh6-Cre:Lmna*^F/F^ cardiomyocytes and subsequently reduced in *Myh6-Cre:Lmna*^F/F^:*Kdm5a*^F/F^ CMs (Fig. 7J) likely reflecting secondary effects associated with phenotypic rescue or indirect regulation through transcription factor–dependent mechanisms. Pathway analysis showed these peaks are localized at inflammatory and extracellular matrix (ECM)-related genes implicated in fibrosis and cell death (Figure 7K and Supplementary Table 2A and B).

A total of 388 H3K4me3 peaks were reciprocally regulated in *Myh6-Cre:Lmna^F/F^:Kdm5a^F/F^*compared to *Myh6-Cre:Lmna^F/F^* (Supplementary Table 2A and 2B). Incorporating the transcript data with the H3K4me3 peaks led to the identification of 189 genes whose transcript levels, as well as H3K4me3 peaks, were rescued. This subset included key cardiac transcription factors TBX5 and members of the ERR family of proteins. The list of loci that showed concordant changes in both transcript levels and promoter H3K4me3 signal in *Myh6-Cre:Lmna*^F/F^ and were normalized upon *Kdm5a* deletion is provided in Supplementary Figure 2 and Supplementary Table 3. Collectively, these findings suggest that KDM5A mediates targeted demethylation of H3K4me3 at promoter regions of genes involved in cardiac function and OXPHOS.

### Suppression of Cardiac and OXPHOS transcription factors TBX5 and ERR in LMNA-CMP

RNA-seq analysis revealed extensive downregulation of cardiac structural genes and FAO, and OXPHOS in LMNA-CMP cardiomyocytes, changes that were restored upon *Kdm5a* deletion. Among the genes that exhibited concordant promoter H3K4me3 alterations (Supplementary Tables 2 and 3), the transcription factors TBX5 and ESRRG emerged as key regulators, suggesting that they further mediate and reinforce KDM5A-driven gene suppression. Supporting this, H3K4me3 enrichment was markedly reduced at the promoters and cis-regulatory elements of *Tbx5* (Figure 8A) in LMNA-CMP cardiomyocytes and partially restored following *Kdm5a* deletion (Figure 8A). Consistent with these epigenetic changes, *Tbx5* transcript levels were markedly reduced in LMNA-CMP hearts and restored upon *Kdm5a* deletion, as confirmed by qRT-PCR (Figure 8B). Likewise, downstream TBX5 target genes, including *Irx4*, *Hopx*, *Myl2*, *Myl3*, *Tnni3*, *Tnnt2*, *Myo5c*, *Tcap*, and several potassium channel genes were significantly suppressed in LMNA-CMP myocytes and re-expressed in *Kdm5a*-deficient hearts, as shown by qRT-PCR (Figure 8C). Likewise, H3K4me3 and RNA levels for another TF, namely *Esrrg* showed reduction in LMNA-CMP and rescue upon *Kdm5a* deletion (Figure 8D-E). ESRRG-regulated targets including *Esrra* and genes within the OXPHOS and FAO pathways, together with genes involved in mitochondrial translation and ribosome biogenesis, were consistently downregulated in LMNA-CMP hearts and rescued in *Kdm5a*-deficient cardiomyocytes (Figures 8F-8G, Supplementary Figure 3). These findings are consistent with our prior data in iPSC-derived cardiomyocytes, where inhibition of KDM5 led to increased expression of *Tbx5, Esrra, and Esrrg.* Together, these data indicate that KDM5A is associated with suppression of cardiac structural and mitochondrial metabolic programs, in part through promoter-proximal H3K4me3 remodeling and in part through transcriptional networks involving TBX5 and ESRRG.

**Figure 8.**
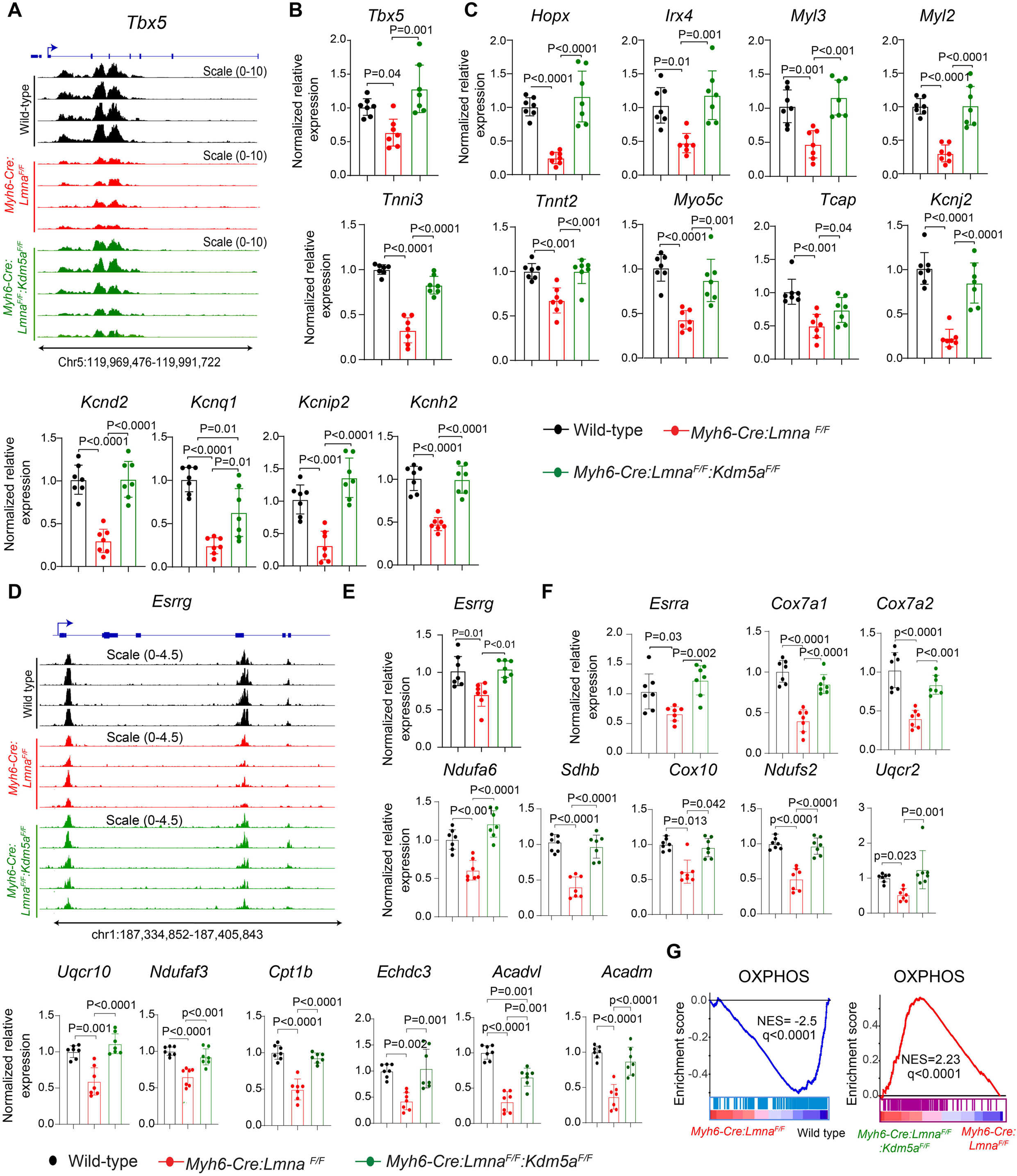
Deletion of *Kdm5a* restores expression of cardiac and OXPHOS transcription factors and their genes in LMNA-CMP. **(A)** Representative IGV genome browser tracks showing H3K4me3 enrichment at cardiac transcription factor *Tbx5* cis regulatory elements *and* promoter region. Tracks shown on mm39 genome assembly. Signals from WT (black), *Myh6-Cre:Lmna^F^*^/F^ (red), and *Myh6-Cre:Lmna^F^*^/F^:*Kdm5a*^F/F^ (green) hearts are displayed. **(B)** Quantitative RT-PCR analysis of *Tbx5* transcript levels across genotypes (n = 7 per group), demonstrating suppression in *Myh6-Cre:Lmna^F^*^/F^ and rescue upon *Kdm5a* deletion. **(C)** Quantitative RT-PCR analysis of downstream targets of TBX5 (n = 7 per group), demonstrating restoration of expression in *Myh6-Cre:Lmna^F^*^/F^:*Kdm5a*^F/F^.**(D)** Representative IGV genome browser tracks showing H3K4me3 enrichment at transcription factor *Esrrg* cis regulatory and promoter region. Signals from Wild-type (black), *Myh6-Cre:Lmna^F^*^/F^ (red), and *Myh6-Cre:Lmna^F^*^/F^:*Kdm5a*^F/F^ (green) hearts are displayed. **(E)** Quantitative RT-PCR analysis of *Esrrg* transcript levels across genotypes (n = 7 per group), demonstrating suppression in *Myh6-Cre:Lmna^F^*^/F^ and rescue upon *Kdm5a* deletion. **(F-G)** Quantitative RT-PCR analysis of ESRRG target genes encoding OXPHOS complex (*Cox7a1, Cox7a2, Ndufa6, Ndufa3, Sdhb, Cox10, Ndufs2, Uqcr2, Uqcr10*) and fatty acid oxidation regulators (*Cpt1b, Acadvl, Acadm, Echdc3*) in *Myh6-Cre:Lmna^F^*^/F^, all of which were significantly restored upon *Kdm5a* deletion (n=6-7 per group). Gene set enrichment analysis demonstrating overall suppression of fatty acid metabolism and OXPHOS in *Myh6-Cre:Lmna^F^*^/F^ compared to Wild-type and partially restored in *Myh6-Cre:Lmna^F^*^/F^:*Kdm5a*^F/F^. Statistical significance was determined by one-way ANOVA and Tukey post hoc pairwise comparisons.

## Discussion

Heart failure is a complex and multisystem disorder leading to cardiac dysfunction and representing the major cause of morbidity and mortality^1,2^. Irrespective of the initiating cause, heart failure progression is driven by convergent molecular derangements encompassing impaired mitochondrial oxidative capacity, disrupted sarcomere integrity, and widespread metabolic reprogramming^3,7,10,26–28^. In this work, we demonstrate KDM5A is upregulated in cardiomyocytes in a mouse model of HF (*Myh6-Cre:Lmna*^F/F^), coinciding with repression of genes required for cardiac contractility and OXPHOS. These changes likely contribute to impaired bioenergetics, cell death, and progressive cardiac dysfunction. We further show that genetic deletion of the histone demethylase KDM5A is sufficient to attenuate cardiac and OXPHOS gene dysregulation and lead to prolonged survival, improved cardiac function, attenuated myocardial cell death and reduced fibrosis in a mouse model of HF. Thus, our findings highlight KDM5A as an important mediator of LMNA-CMP and HF phenotype and as a potential epigenetic therapeutic target.

Our study also identified key cardiac transcription factors, namely TBX5 as well as mitochondrial regulators ESRRG/ ESRRA, as targets of KDM5A among others. The roles of these factors in heart failure and especially in cardiac gene expression and mitochondrial function are well established ^29–32^. Consistent with this, our previous data in iPSC-CMs showed that inhibition of KDM5 increased H3K4me3 deposition at the ESRRA promoter, accompanied by elevated RNA and protein levels and knockdown of ESRRA in KDM5 inhibited suppression of expression of a subset of KDM5 target genes in iPSC-CMs, providing further evidence that ERR proteins are direct targets of KDM5^22^. Importantly, our current findings extend this paradigm by uncovering the regulation of TBX5, cardiac transcription factors not previously linked to KDM5A activity.

Our findings uncover a previously unrecognized role of KDM5A in the pathogenesis of LMNA-CMP. Genome-wide profiling revealed a global reduction of H3K4me3 in LMNA-deficient cardiomyocytes, despite the loss of lamin-associated domains (LADs), which are typically associated with transcriptional repression^19,33,34^. This suggests that KDM5A activation establishes a unique repressive state in LMNA-deficient cardiomyocytes. Interestingly, attenuation of LMNA cardiomyopathy severity has also been reported following BET bromodomain inhibition with JQ1, which similarly targets transcriptional programs as an epigenetic reader^9,35^. Although mechanistically distinct, the BET proteins excerbate pathological programs by inducing transcriptional elongation while the KDM5A seems to suppress cardiac and OXPHOS gene programs, however, both BET inhibition and KDM5A deletion converge on restoring cardiac function, suggesting that modulation of epigenetic readers and erasers represents a shared vulnerability in LMNA-driven cardiomyopathy and HF. Whether combined targeting of these pathways would yield additive benefit remains an important question for future investigation.

The KDM5 family of histone demethylases regulates transcription by removing methyl groups from H3K4me3/2/1 and thereby suppressing expression. However, KDM5A can also activate or modulate gene expression by interacting with co-factors and cell-type-specific transcription factors^18,36–39^. In this study, while several genes exhibited direct promoter-associated H3K4me3 changes in a KDM5A-dependent manner, a distinct subset showed restored expression upon *Kdm5a* deletion without detectable changes in H3K4me3 deposits in their promoters. The former likely represent direct KDM5A targets, whereas the latter reflect downstream effects mediated through transcription factors such as TBX5, ESRRA, and ESRRG. We also observed that LMNA-CMP hearts like most forms of HF displayed upregulation of inflammatory and epithelial-to-mesenchymal transition (EMT)–related gene programs. These changes may be attributable to either indirect effects as noted above or a result of KDM5A’s function within multiprotein complexes, like retinoblastoma protein (RB1), Sin3B/NuRD complexes, and transcription factors including E2Fs and ZBTB20 ^36,38,39^. Such scaffolding interactions may also explain why *Kdm5a* deletion normalizes transcription even in the absence of measurable promoter H3K4me3 changes. In addition, KDM5A may contribute to disease progression by directly engaging pro-inflammatory transcription factors, repressing anti-inflammatory regulators, or indirectly activating stress-responsive pathways downstream of mitochondrial dysfunction and cell death. Collectively, these findings highlight KDM5A’s broad regulatory influence across multiple transcriptional networks in the failing heart and establish it as a driver of HF pathogenesis.

The LMNA-CMP model provides a severe and well-defined genetic model of cardiomyopathy to interrogate the epigenetic mechanisms driving heart failure progression. Although complete loss of LMNA represents a severe phenotype, this model demonstrates the key pathological features seen in heterozygous LMNA deficiency and disease-causing LMNA mutations, including cardiac dysfunction, apoptosis, fibrosis, and mitochondrial abnormalities ^9,40,41^. Because disease progression is rapid and severe, this model allows mechanistic insights to be obtained acutely. In this context, a partial improvement in cardiac function and survival are meaningful, given the severe nature of the disease and the multiple pathways involved in heart failure. Importantly, interventions that show benefit in this setting are likely to have equal or greater impact in milder, clinically relevant LMNA cardiomyopathy models.

Although the precise mechanism underlying KDM5A induction in CMs remains unknown, it is noteworthy that KDM5A has been identified as an interacting partner of RB1^42–44^. Notably, RB1 expression is regulated by its interaction with LMNA^45,46^. Therefore, in LMNA-CMP and in heart failure, the loss of RB1 may stabilize KDM5A and facilitate its relocalization to gene promoters, thereby providing a plausible mechanism for its aberrant activation. Mice with the deletion of *Kdm5a* alone were not included in the mechanistic studies, because they did not show a discernible phenotype. Nevertheless, the possibility that some of the transcriptional or chromatin effects reflect baseline consequences of *Kdm5a* loss rather than disease-specific reversal could not be excluded.

Together, these findings implicate KDM5A as a key epigenetic regulator orchestrating pathological transcriptional programs in HF, encompassing genes governing mitochondrial metabolism, contractile function, and OXPHOS. Notably, systemic or cardiomyocyte-specific deletion of *Kdm5a* was well tolerated in adult mice, underscoring the therapeutic viability of its inhibition ^37^. Thus, by revealing KDM5A as a driver of disease progression, this study highlights a rational avenue for intervention, supported by the growing repertoire of selective KDM5 inhibitors and RNAi-based strategies capable of mitigating pathological remodelling in LMNA-CMP and broader heart failure contexts.

## Methods

### Anesthesia and Euthanasia

All echocardiography and tissue isolation procedures were conducted under general anesthesia, induced with 3% inhaled isoflurane and maintained at 0.5–1.0% throughout^9,20,21,25^. For cardiomyocyte (CM) and whole heart (WH) isolations, mice were euthanized by cervical dislocation while under anesthesia.

### Mice Used in the Study

Wild-type (C57BL/6J), *Myh6-Cre* (C57BL/6J), and *Kdm5a* floxed mice (JB6.129S6(Cg)-Kdm5a^tm1Kael^/J) were obtained from Jackson Laboratory. *Lmna*^F/F^ and *Kdm5a*^F/F^ mouse lines were backcrossed to C57BL/6J wild-type mice to homogenize the genetic background. Cardiac-specific *Lmna* knockout mice (*Myh6-Cre:Lmna*^F/F^) were generated by crossing *Lmna*^F/F^ mice with *Myh6-Cre* transgenic mice. *Myh6-Cre:Kdm5a*^F/F^ mice were generated by breeding *Myh6-Cre* mice with *Kdm5a*^F/F^ mice and were subsequently crossed with *Lmna*^F/F^ mice to generate cardiac-specific double-knockout mice (*Myh6-Cre:Lmna*^F/F^:*Kdm5a*^F/F^). Genotyping was performed by PCR using DNA extracted from mouse tails, following incubation in buffer at 55°C for 15 minutes and 95°C for 3 minutes. PCR was conducted with 2 μL of extract using a ready-mix blue extract. Oligonucleotide primers are listed in Supplementary Table 4.

### Sex as a biological variable

Age-matched male and female mice were included in all experimental groups. Data were analyzed with sex incorporated as a biological variable to evaluate potential sex-dependent differences in the observed phenotypic outcomes.

### Survival Analysis

Kaplan–Meier survival curves were generated for each group, using approximately equal numbers of male and female mice.

### Echocardiography

Cardiac function was assessed using a Vevo 1100 system (Fujifilm VisualSonics) under 1% isoflurane anesthesia, maintaining a heart rate of ~500 bpm. M-mode images were analyzed to measure interventricular septal thickness (IVST), posterior wall thickness (PWT), left ventricular end-diastolic diameter (LVEDD), and end-systolic diameter (LVESD) over 3-4 cardiac cycles^9,20,25^. To minimize bias, echocardiographic measurements were performed by investigators blinded to experimental group allocation.

### Cardiomyocyte Isolation

Cardiomyocytes were isolated from P16 and P21 mice following published protocols ^9,20,25,47^. Briefly, hearts were mounted on a Langendorff system and perfused with collagenase (1.2 mg/mL, Worthington cat # LS004176) at 4 mL/min for 10–12 minutes. Ventricles were minced in a stop buffer (10% calf serum, 12.5 µM CaCl_₂_, 2 mM ATP), filtered (100 µm), and centrifuged at 300 rpm for 4 minutes. The CM pellet was resuspended in appropriate buffers for downstream applications.

### Immunoblotting

Protein expression in whole heart and cardiac myocyte (CM) extracts was analyzed by immunoblotting (IB). Tissues and or CM were homogenized in RIPA buffer containing benzonase (250units/ml buffer) with protease/phosphatase inhibitors (Roche #4693116001 and #49068459001). Lysates were nutated at 4°C for 30 minutes and sonicated (3 cycles of 30 s ON/OFF). Supernatants were collected after centrifugation (14,000 rpm, 15 minutes, 4°C). Extracts (30–40 µg) were resolved by SDS-PAGE and transferred to PVDF membranes at 4°C overnight. Antibodies used in the manuscript are listed in Supplementary Table 4.

### Quantitative Real-time PCR (qPCR)

Gene expression was quantified by qPCR using gene-specific primers and SYBR Green master mix, normalized to GAPDH. Primer sequences are in Supplementary Table 4.

### RNA Sequencing

RNA-seq was performed from P18–P20 isolated CMs. RNA was extracted with the RNeasy Mini Kit; samples with RIN >8.7 were used. rRNA was removed, and libraries were generated for 150 bp paired-end sequencing (Illumina NextSeq500, Genewiz). Read alignment was completed using the Pluto Bioinformatics platform. DEGs (q-value <0.05) were obtained using limma-voom.

### Cleavage Under Targets and Release Using Nuclease (CUT&RUN) Assay

CUT&RUN was performed by the Epigenomics Profiling Core at MD Anderson Cancer Center with some modifications^48^. Briefly, cardiomyocytes (CMs) isolated from P18–P20 mice were mildly crosslinked with 0.1% formaldehyde for 1 minute and quenched with 125 mM glycine. Nuclei isolated from ~500,000 cells were immobilized on Concanavalin A-coated magnetic beads (Bangs Laboratories) and permeabilized using wash buffer containing 0.02% digitonin (Promega). Bead-bound nuclei were incubated with rabbit IgG (Millipore) and H3K4me3 (EpiCypher) antibodies overnight at 4°C. The next day, targeted chromatin digestion was achieved by pAG-MNase (EpiCypher) binding and incubation with CaCl_2_ at 4°C followed by purification of DNA using MinElute columns (Qiagen). Library preparation was carried out using the NEBNext Ultra II DNA Library Prep Kit (New England Biolabs) according to the manufacturer’s instructions, and sequencing was performed on an Illumina NextSeq 500 platform to obtain 15–20 million reads per sample.

Sequencing reads were aligned to the mouse reference genome (GRCm39), and BAM files were used for peak calling with MACS2 using standard parameters (effective genome size = 1.87e+09, band width = 300, model fold = [10, 30], and q value cutoff = 5.00e-02). Peak quality assessment and enrichment profiling were conducted using the computeMatrix and plotHeatmap functions in deepTools. Heatmaps were generated from log-transformed normalized values [log(CPM+1)]. Differential peak analysis between genotypes Wild-type (WT) vs. *Myh6-Cre:Lmna^F/F^* and *Myh6-Cre:Lmna^F/F^:Kdm5a^F/F^*vs. *Myh6-Cre:Lmna^F/F^* was performed using DiffBind. Peaks were annotated and their genomic distribution was analyzed using Chip seeker, and visualization was performed with deepTools. Read alignment was completed using the Pluto Bioinformatics platform, and all downstream analyses, including peak calling and differential analysis, were conducted on a local Galaxy instance or in R.

### Pathway Analysis

Gene set enrichment analysis (GSEA) was performed on normalized counts using MSigDB gene sets, with 1,000 permutations and q<0.05, or using the enrich R online tool.

### Apoptosis

Hearts from P21 mice were fixed in 10% formalin overnight, dehydrated through ethanol gradients, and embedded in paraffin^9,20,25,47,49–51^. Sections (5 µm) were placed on Superfrost slides. TUNEL assays were conducted using the In-Situ Cell Death Detection Kit (Roche # 11684795910) following deparaffinization and proteinase K digestion (20 µg/mL, 20 minutes, RT). Nuclei were counterstained with 1 µg/mL DAPI. Images were captured with a Zeiss Axioplan microscope and quantified in ImageJ from 40X high-magnification fields (6–9 sections/heart, ≥5 mice/group). Quantitative analysis was performed in a blinded manner.

### Fibrosis

Sections were stained with Picrosirius red for 1 hour, washed in ethanol, and mounted^9,20,25,47,49–51^. Collagen volume fraction (CVF) was analyzed in 10 fields/6 sections/5 mice/group using ImageJ. Quantitative analysis was performed in a blinded manner.

### Statistics

For genomic studies, differential gene expression and peak enrichment analyses were performed using a false discovery rate (FDR) threshold of 5% (*q* < 0.05). Pathway enrichment analyses were conducted with Ingenuity Pathway Analysis (IPA), applying a *z*-score cutoff of ±2 and a significance threshold of p< 0.05. For other studies, the normality of continuous variables was assessed using the Shapiro–Wilk test. Normally distributed variables were compared between two groups using the two-sample *t*-test and across multiple groups using one-way ANOVA followed by Tukey’s pairwise comparison. Non-normally distributed variables were compared using the Mann–Whitney test between two groups and by the Kruskal-Wallis test followed by Dunn’s test for multiple comparisons across groups. Categorical variables were analyzed using the Chi-square test. Survival analyses were performed using Kaplan–Meier survival curves. For molecular and histological analyses, one mouse per litter was randomly selected to preserve statistical independence. Survival and echocardiographic assessments included multiple animals from the same litter but selected randomly data were analyzed assuming independence between observations. Survival was compared using the log-rank (Mantel–Cox) test. In Kaplan–Meier curves, stepwise declines indicate deaths and symbols denote censored animals alive at the time of analysis. Mice were euthanized at humane endpoints defined by institutional veterinary staff or died spontaneously. All statistical analyses were performed using R and/or GraphPad Prism version 7.

## Supporting information

uncropped WBs

## Study Approval

All animal experiments were conducted following the NIH Guidelines for the Care and Use of Laboratory Animals and were approved by the Institutional Animal Care and Use Committee at the University of Texas Health Science Center at Houston (AWC-24-0021).

## Data Availability

RNA-seq and CUT&RUN data have been deposited in [GSE309726] and additional processed data are presented in Supplementary Table 1-3.

## Funding support

PG is supported by the NIH, NHLBI, award R01HL174481. AJM is supported by the NIH, NHLBI Awards: R01HL151737 and R01HL132401 and NIA Award: R01AG082751. Dr. Rouhi is supported by the American Heart Association Career Development Award 23CDA1053101. This work was also supported by John. Dunn Foundation (P.G.) and the Ewing Halsell Foundation (A.J.M).

## Notes

### Competing Interest Statement

The authors have declared no competing interest.

