## Supplementary figures and images for "The Epigenetic Regulator Histone Demethylase KDM5A is Activated and Pathogenic in a Mouse Model of Heart Failure"

### Supplementary Figure 1

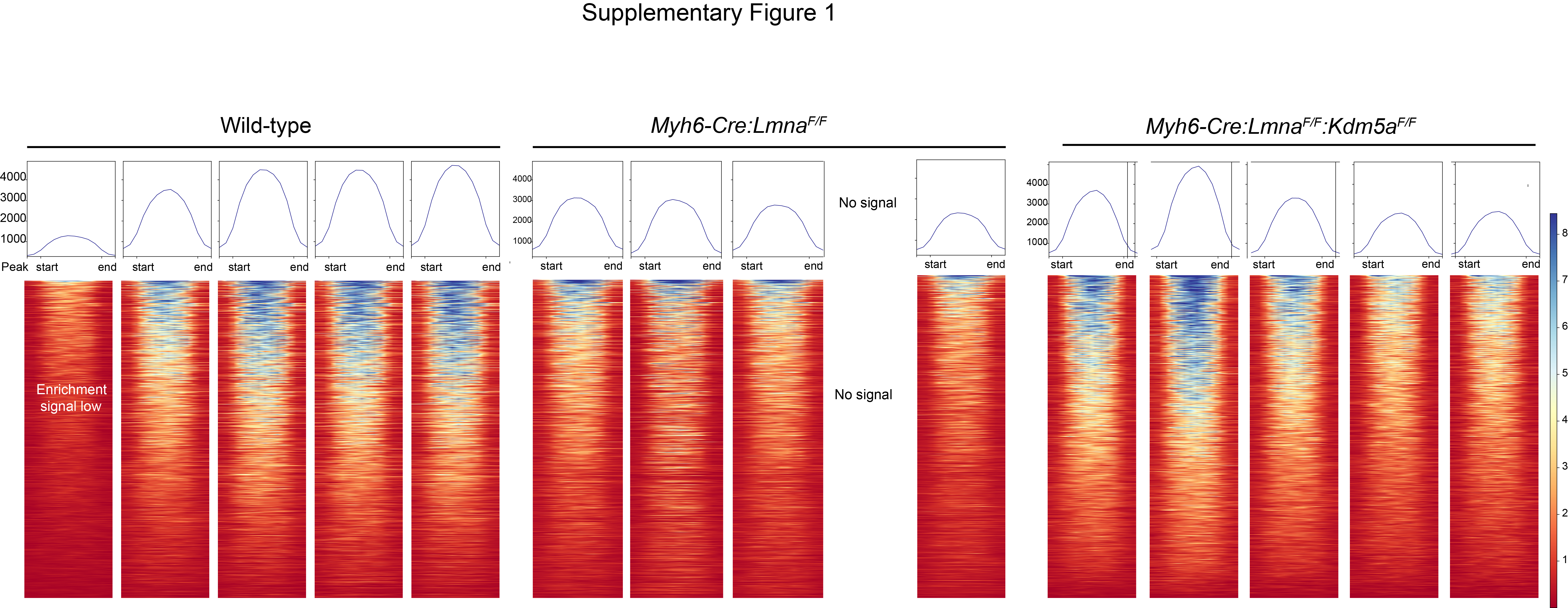

### Supplementary Figure 2

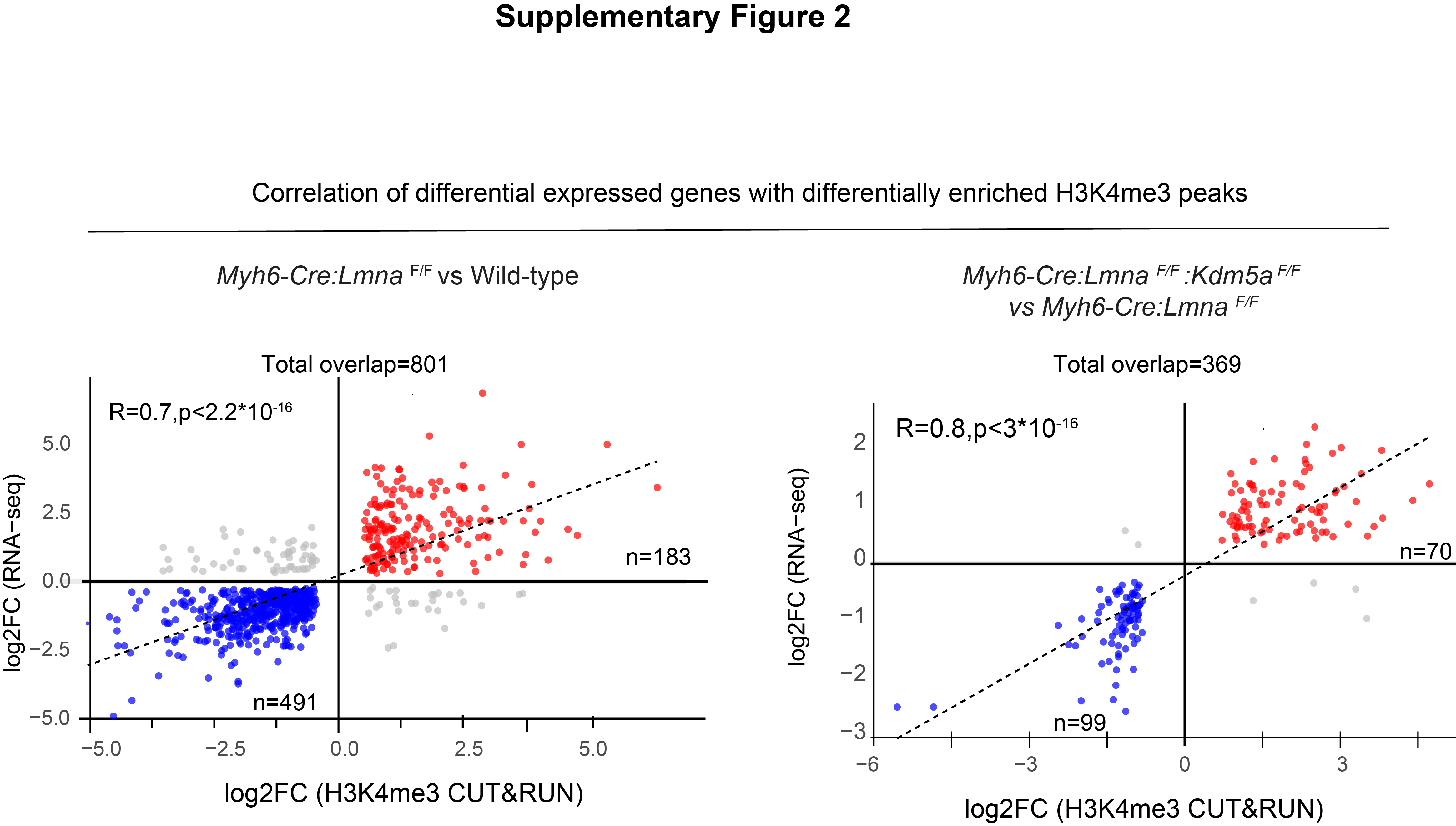

### Supplementary Figure 3

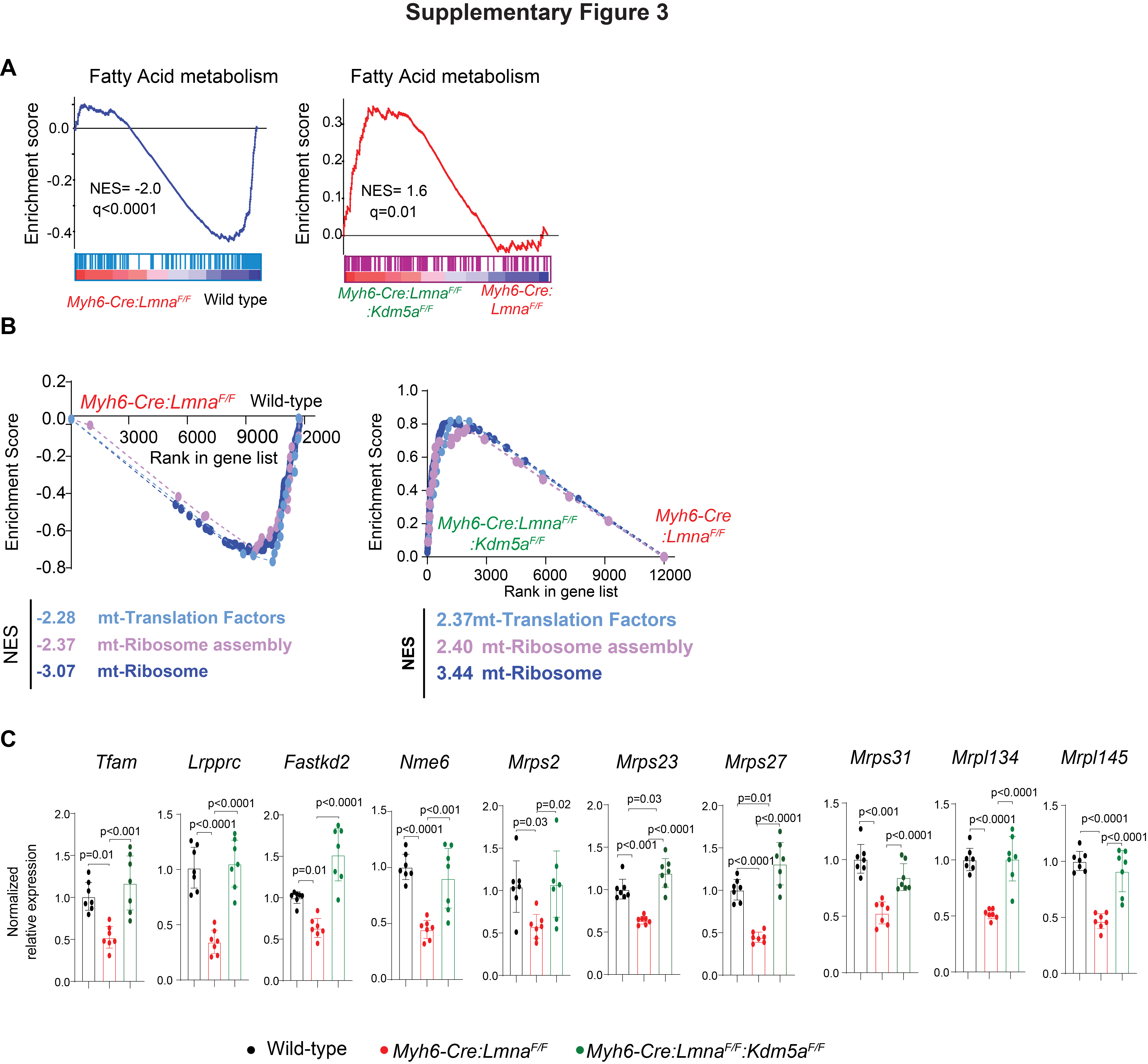
