## Supplementary Figure Legend for "The Epigenetic Regulator Histone Demethylase KDM5A is Activated and Pathogenic in a Mouse Model of Heart Failure"

**Supplementary Figure 1: Quality control analysis of CUT&RUN H3K4me3 data**

**(A)** Quality control assessment showing heatmaps of H3K4me3 enrichment at transcription start sites (±3 kb) from CUT&RUN experiments in cardiomyocytes from WT, *Myh6-Cre:Lmna*^F/F^, and *Myh6-Cre:Lmna*^F/F^:*Kdm5a*^F/F^ CM. Robust enrichment was observed in all WT samples except WT1, and one *Myh6-Cre:Lmna*^F/F^ sample showed no detectable peaks, likely reflecting failed experiments. These two samples were excluded from downstream analyses.

**Supplementary Figure 2: Correlation between H3K4me3 enrichment and gene expression.**

**(A)** Correlation analysis of H3K4me3 enrichment and RNA expression in WT versus *Myh6-Cre:Lmna*^F/F^ cardiomyocytes. A total of 801 differential H3K4me3 peaks overlapped with annotated genes. Of these, 183 genes showed concordant upregulation at both the RNA level and promoter-associated H3K4me3, while 491 genes exhibited concordant downregulation. The Pearson correlation coefficient and corresponding p values for the 801 genes are indicated.

**(B)** Similar analysis comparing *Myh6-Cre:Lmna*^F/F^:*Kdm5a*^F/F^ versus *Myh6-Cre:Lmna*^F/F^ cardiomyocytes. Here, 369 differential H3K4me3 peaks overlapped with annotated genes, including 70 showing concordant upregulation and 99 showing concordant downregulation at both RNA and H3K4me3 levels.

**Supplementary Figure 3: Rescue of ERR targets expression upon *Kdm5a* deletion in *Myh6-Cre:Lmna*^F/F^** **CM.**

**(A-B)** GSEA plots from RNA-seq analysis showing genes belonging to FAO, mitochondrial translation and ribosome assembly that are targets of ERR family of TF showing suppression in *Myh6-Cre:Lmna*^F/F^ CM and restoration upon *Kdm5a* deletion in *Myh6-Cre:Lmna*^F/F^:*Kdm5a*^F/F^.

**(C)** Quantitative RT-PCR analysis from independent samples showing genes involved in mitochondrial DNA synthesis and maintenance (*Tfam, Polrmt, Tf2bm*) and RNA processing (*Nme6, Lrrpc*), and translation are suppressed in *Myh6-Cre:Lmna*^F/F^ CM and restoration upon *Kdm5a* deletion in *Myh6-Cre:Lmna*^F/F^:*Kdm5a*^F/F^ CMs.
