## Supplementary Table 4 for "The Epigenetic Regulator Histone Demethylase KDM5A is Activated and Pathogenic in a Mouse Model of Heart Failure"

**Antibodies**

| Primary Antibody | Primary Source | Catalog No. | Company | Dilution |
| --- | --- | --- | --- | --- |
| KDM5A | Rabbit | Custom,KDM5A  (Region 1311-1359) | Pacific Immunology | 1:1000 in milk |
| GAPDH PolyAb | Rabbit | 2118S | Cell Signaling | 1:1000 in milk |
| GAPDH McAb | Mouse | 97166S | Cell Signaling | 1:1000 in milk |
| Vinculin McAb | Mouse | 66305-2-Ig | Proteintech | 1:1000 in BSA |
| mAb to Vinculin [EPR8185] | Rabbit | ab129002 | Abcam | 1:1000 in BSA |
| LMNA | Rabbit | ab26300 | Abcam | 1:1000 in BSA |
| Histone H3 (D1H2) | Rabbit | 4499S | Cell Signaling | 1:1000 in BSA |
| Caspase-3 | Rabbit | 9662 | Cell Signaling | 1:1000 in BSA |
| Bad | Rabbit | 9292T | Cell Signaling | 1:1000 in BSA |
| BAX PolyAb | Rabbit | 50599-2-Ig | Proteintech | 1:1000 in BSA |

| Secondary Antibody | Primary Source | Catalog No. | Company | Dilution |
| --- | --- | --- | --- | --- |
| Anti-rabbit IgG, HRP | Goat | 7074 | Cell Signaling | 1:3000 in milk |
| Anti-mouse IgG, HRP | Horse | 7076 | Cell Signaling | 1:3000 in milk |

**Primers used for Genotyping**

| **Genes** | **Primer Sequence 5’-3’** |
| --- | --- |
| Lmna-F | TCGAGGCTCTTCTCAACTCC |
| Lmna-R | CTCTCCTCTGAAGTGCTTGGA |
| Cre-F | ATGACAGACAGATCCCTCCTATCTCC |
| Cre-R | CTCATCACTCGTTGCATCATCGAC |
| Cre-Control-F | CAAATGTTGCTTGTCTGGTG |
| Cre-Control-R | TCAGTCGAGTGCACAGTTT |
| Kdm5a-F | GTTTGAATTTCACTCTATGCTGGG |
| Kdm5a-R | TCCCAGCATGGATCTTGTCC |

**Primers used for QPCRs**

| **Genes** | **Primer Sequence 5’-3’** |
| --- | --- |
| Mmu-Acadm-F | AACACAACACTCGAAAGCGG |
| Mmu-Acadm-R | TTCTGCTGTTCCGTCAACTCA |
| Mmu-Acads-F | GACTGGCGACGGTTACACA |
| Mmu-Acads-R | GGCAAAGTCACGGCATGTC |
| Mmu-Acadvl-F | ACTACTGTGCTTCAGGGACAA |
| Mmu-Acadvl-R | GCAAAGGACTTCGATTCTGCC |
| Mmu-Acta1-F | CCCAAAGCTAACCGGGAGAAG |
| Mmu-Acta1-R | GACAGCACCGCCTGGATAG |
| Mmu-Anxa1-F | ATGGGGAATCGTCATGCCAAA |
| Mmu-Anxa1-R | CTTCTGCTTTATCTGTTGCCTCT |
| Mmu-Atp2a2-F | TCACACAAAGACCGTGGAGG |
| Mmu-Atp2a2-R | TCTTCAGCCGGCAATTCGTT |
| Mmu-Bcl2l2-F | GCGGAGTTCACAGCTCTATAC |
| Mmu-Bcl2l2-R | AAAAGGCCCCTACAGTTACCA |
| Mmu-Bid-F | GCCGAGCACATCACAGACC |
| Mmu-Bid-R | TGGCAATGTTGTGGATGATTTCT |
| Mmu-Caspase3-F | TGGTGATGAAGGGGTCATTTATG |
| Mmu-Caspase3-R | TTCGGCTTTCCAGTCAGACTC |
| Mmu-Cd44-F | GGCTCTGATTCTTGCCGTCT |
| Mmu-Cd44-R | TCCTGTCTTCCACTGTCCCA |
| Mmu-Col1A1-F | TGACTGGAAGAGCGGAGAGTA |
| Mmu-Col1A1-R | GAGTAGGGAACACACAGGTCT |
| Mmu-Col3a1-F | CTGTAACATGGAAACTGGGGAAA |
| Mmu-Col3a1-R | CCATAGCTGAACTGAAAACCACC |
| Mmu-Col5A2-F | GTACCACTGGGCAAAGAGGA |
| Mmu-Col5A2-R | CTTTTCCTGGTGTACCCGCT |
| Mmu-Col16a1-F | CCAGGGCATCTTGGTCCTC |
| Mmu-Col16a1-R | CCGCCAAGCCCCTTTCTC |
| Mmu-Cox10-F | AGAAGAGCTATACAGGGATTGCC |
| Mmu-Cox10-R | CTGTGTGACATACATGCGCTT |
| Mmu-Cox7a1-F | GCTCTGGTCCGGTCTTTTAGC |
| Mmu-Cox7a1-R | GTACTGGGAGGTCATTGTCGG |
| Mmu-Cox7a2-F | GCTGGCCCTTCGTCAGATT |
| Mmu-Cox7a2-F | GCTGGCCCTTCGTCAGATT |
| Mmu-Cpt1a-F | TGGCATCATCACTGGTGTGTT |
| Mmu-Cpt1a-R | GTCTAGGGTCCGATTGATCTTTG |
| Mmu-Cpt1b-F | GACTTCCGGCTTAGGCGGG |
| Mmu-Cpt1b-R | GAATAAGGCGTTTCTTCCAGGA |
| Mmu-Ctgf-F | AACCGCAAGATCGGAGTGT |
| Mmu-Ctgf-R | CTGCGGTACACCGACCCAC |
| Mmu-Echdc3-F | CGACAGCAGGACGGAATCAG |
| Mmu-Echdc3-R | GTGAAGAATGTCACTTCGGAGAG |
| Mmu-Ehhadh-F | ATGGCTGAGTATCTGAGGCTG |
| Mmu-Ehhadh-R | ACCGTATGGTCCAAACTAGCTT |
| Mmu-Esrra-F | GGGGAGCATCGAGTACAGC |
| Mmu-Esrra-R | AGACGCACACCCTCCTTGA |
| Mmu-Esrrg-F | GCCATCAGAACGGACTGGAC |
| Mmu-Esrrg-R | GCGATGTCGCCACACACTA |
| Mmu-Fastkd2-F | ATGAATAGCAAAGCACGTTCCT |
| Mmu-Fastkd2-R | AGTTCCCAATGGATCTATCCTCA |
| Mmu-Gadd45a-F | AGACCGAAAGGATGGACACG |
| Mmu-Gadd45a-R | GTACACGCCGACCGTAATG |
| Mmu-Gadd45b-F | GAGGCGGCCAAACTGATGAAT |
| Mmu-Gadd45b-R | CGCAGCAGAACGACTGGAT |
| Mmu-Gapdh-R | TGGCAAAGTGGAGATTGTTGC |
| Mmu-Gapdh-R | AAGATGGTGATGGGCTTCCCG |
| Mmu-Hopx-F | GGGCTGAGTGAAAAGTCCCA |
| Mmu-Hopx-R | CACTGAAAGTCGCCTGACCT |
| Mmu-Irx4-F | TCCTACCCGCAGTTTGGATAC |
| Mmu-Irx4-R | GCGGCAGAATTGAGTTCGT |
| Mmu-Kcnd2-F1 | TCAGGACGCTCTGATAGTGCT |
| Mmu-Kcnd2-R1 | TCTGGGTATCGTTCCAGGGTG |
| Mmu-Kcnh2-F1 | GTGCTGCCTGAGTATAAGCTG |
| Mmu-Kcnh2-R1 | CCGAGTACGGTGTGAAGACT |
| Mmu-Kcnip2-F | GGATGAGTTTGAACTATCCACGG |
| Mmu-Kcnip2-R | GACAATTCCGCTGGGACATTC |
| Mmu-Kcnj2-F | AACCGCTACAGCATCGTCTC |
| Mmu-Kcnj2-R | GTTGTCGGGTATGGACTTTACTC |
| Mmu-Kcnq1-F1 | CGCGGTGGTCAAGAAGTGT |
| Mmu-Kcnq1-R1 | GCACTGTAGATGGAGACCCG |
| Mmu-Kdm5a-F | GGTCTCTTTTGAAGTCACAC |
| Mmu-Kdm5a-R | CTTCACAGGCAAATGGAGGTT |
| Mmu-Lgals3-F | AGACAGCTTTTCGCTTAACGA |
| Mmu-lgals3-R | GGGTAGGCACTAGGAGGAGC |
| Mmu-Lmna-F | GGCCCGGCTCAAGGACC |
| Mmu-Lmna-R | TCTCCCAGGGCCGCCTC |
| MMu-Lrpprc-F | TCTGGGACAAACTTCAGCAGT |
| MMu-Lrpprc-R | TGATCGGAAGGTCTTTCGTTTTC |
| Mmu-Mhy6-F | GCCCAGTACCTCCGAAAGTC |
| Mmu-Mhy6-R | ATCAGGCACGAAGCACTCC |
| Mmu-Mhy7-F | ACTGTCAACACTAAGAGGGTCA |
| Mmu-Mhy7-R | TTGGATGATTTGATCTTCCAGGG |
| Mmu-Mmp9-F | GGACCCGAAGCGGACATTG |
| Mmu-Mmp9-R | CGTCGTCGAAATGGGCATCT |
| Mmu-Mrpl134-F | TCGGTGGCCCTATTGAGTG |
| Mmu-Mrpl134-R | GCTCGGCTGATACTCGTTTCC |
| Mmu-Mrpl45-F | AGACCTTGATGATTCCTGGTCC |
| Mmu-Mrpl45-R | CCCTTGAGTCTCTTCGTACTCC |
| Mmu-Mrps23-F | GGCCGGGGTGTTGAAAGAG |
| Mmu-Mrps23-R | CTTTGCCGTATCGCAAGCG |
| Mmu-Mrps27-F | GCCTTTGCCTCGCAGGTAA |
| Mmu-Mrps27-R | TGTCCACAAACCGTGATATTGAT |
| Mmu-Mrps2-F | ATCCTAGATACGCCATTACAGCA |
| Mmu-Mrps-R | GGCGGCAACCAGCTTTATG |
| Mmu-Mrps31-F | CTCCACAGAATCCCGGCATTT |
| Mmu-Mrps31-R | ACTGGTCAACTTTCTTGCTACAG |
| Mmu-Mt-Atp6-F | TAGCCCACCAACAGCTACCA |
| Mmu-Mt-Atp6-R | GGAGGGTGAATACGTAGGCT |
| Mmu-Mt-Cox1-F | GCTAGCCGCAGGCATTACTA |
| Mmu-Mt-Cox1-R | CTCCTCCAGCGGGATCAAAG |
| Mmu-Mt-Cox2-F | AACCGAGTCGTTCTGCCAAT |
| Mmu-Mt-Cox2-R | CTAGGGAGGGGACTGCTCAT |
| Mmu-Mt-Cox3-F | GGCCACCACACTCCTATTGT |
| Mmu-Mt-Cox3-R | ACGCTCAGAAGAATCCTGCAA |
| Mmu-Mt-Nd1-F | TCCGAGCATCTTATCCACGC |
| Mmu-Mt-Nd1-R | GTATGGTGGTACTCCCGCTG |
| Mmu-Mt-Nd5-F | ATTCGGAAGCATCTTTGCAGG |
| Mmu-Mt-Nd5-R | TGTGAGGACTGGAATGCTGG |
| Mmu-Myl2-F | ATCGACAAGAATGACCTAAGGGA |
| Mmu-Myl2-R | ATTTTTCACGTTCACTCGTCCT |
| Mmu-MYL3-F | TGCCTCCAAGATTAAGATCGAGT |
| Mmu-MYL3-R | CTCTGCCTGGGTAGGATTCTG |
| Mmu-Myo5c-F | AAGTCGGCTGAAATCGCAAAG |
| Mmu-Myo5c-R | CGGATTCTGAGATTATGCAGCAC |
| Mmu-Ndufa5-F | ATGGCGGGCTTGCTGAAAA |
| Mmu-Ndufa5-F | GCTGCATGTTTAGGAAAGTGCTT |
| Mmu-Ndufa6-F | TCGGTGAAGCCCATTTTCAGT |
| Mmu-Ndufa6-R | CTCGGACTTTATCCCGTCCTT |
| Mmu-Ndufaf3-F | CAAAGCGAGTTCCCTCAGG |
| Mmu-Ndufaf3-R | ACACGCGATTTCCGCAGATAG |
| Mmu-NdufB8-F | TGTTGCCGGGGTCATATCCTA |
| Mmu-NdufB8-R | AGCATCGGGTAGTCGCCATA |
| Mmu-Ndufs2-F | ACTGCCATTGAAGCTCCTAAG |
| Mmu-Ndufs2-R | TGCCAACATGTGTCCCTTAG |
| Mmu-Nme6-F | CCTCATTGTACGAACGAGGGA |
| Mmu-Nme6-R | GCAAGGATATAGGCTCGGATTG |
| Mmu-Nppa-F | GTGCGGTGTCCAACACAGAT |
| Mmu-Nppa-R | TCCAATCCTGTCAATCCTACCC |
| Mmu-Nppb-F | GAGGTCACTCCTATCCTCTGG |
| Mmu-Nppb-R | GCCATTTCCTCCGACTTTTCTC |
| Mmu-Opa1-F | GTGTGCTGGAAATGATTGCTC |
| Mmu-Opa1-R | TGGTGAGATCAAATTCCCGAG |
| Mmu-Polrmt-F | TGGGCGCAAAAGCTAGAGG |
| Mmu-Polrmt-R | GTGAAGGGTCCAGAACTCCTG |
| Mmu-Postn-F | AGAGAAATCCCTGCACGACA |
| Mmu-Postn-R | GTTGGTGCAAACAAGGTCCA |
| Mmu-Sdhb-F | AATTTGCCATTTACCGATGGGA |
| Mmu-Sdhb-R | AGCATCCAACACCATAGGTCC |
| Mmu-Sdhd-F | TGGTCAGACCCGCTTATGTG |
| Mmu-Sdhd-R | GGTCCAGTGGAGAGATGCAG |
| Mmu-Sfrp1-F | TACTGGCCCGAGATGCTCAA |
| Mmu-Sfrp1-R | GAGGCTTCCGTGGTATTGGG |
| Mmu-Sfrp2-F | CCCTTTGTAAAAATGACTTCGCAC |
| Mmu-Sfrp2-R | CAGGATGATCTTGGTGTCTCTGT |
| Mmu-Spp1-F | AGCAAGAAACTCTTCCAAGCAA |
| Mmu-Spp1-R | GTGAGATTCGTCAGATTCATCCG |
| Mmu-Tbx5-F | ATGGCCGATACAGATGAGGG |
| Mmu-Tbx5-R | TTCGTGGAACTTCAGCCACAG |
| Mmu-Tcap-F | CGTGGGCTACAGGAATACCAG |
| Mmu-Tcap-R | GAGACATGGATCGAGACAGGG |
| MMu-Tfam-F | AACACCCAGATGCAAAACTTTCA |
| MMu-Tfam-R | GACTTGGAGTTAGCTGCTCTTT |
| Mmu-Tfb2m-F | GGCCCATCTTGCATTCTAGGG |
| Mmu-Tfb2m-R | GCAACGGCTCTATATTGAAGTCA |
| Mmu-Tgfb1-F | TGGAGCAACATGTGGAACTC |
| Mmu-Tgfb1-R | GTCAGCAGCCGGTTACCA |
| Mmu-Tgfb2-F | CTTCGACGTGACAGACGCT |
| Mmu-Tgfb2-R | GCAGGGGCAGTGTAAACTTATT |
| Mmu-Tgfb3-F | AGCTCTTCCAGATACTTCGACC |
| Mmu-Tgfb3-R | AAAGACAGCCATTCAGCGGT |
| Mmu-Tnni3-F | CTCTGCCAACTACCGAGCCTA |
| Mmu-Tnni3-R | CTCTTCTGCCTCTCGTTCCAT |
| Mmu-Tnnt2-F | CAGAGGAGGCCAACGTAGAAG |
| Mmu-Tnnt2-R | CTCCATCGGGGATCTTGGGT |
| Mmu-Uqcr10-F | ATCCCTTCGCGCCTGTACT |
| Mmu-Uqcr10-R | GTGCTCGTAGATCGCGTCT |
| Mmu-Uqcr2-F | AAAGTTGCCCCGAAGGTTAAA |
| Mmu-Uqcr2-R | GAGCATAGTTTTCCAGAGAAGCA |
| Mmu-Uqcrh-F | GTGGACCCCCTAACAACAGTG |
| Mmu-Uqcrh-R | CGGGAAGACACGCGATTATCA |
| Mmu-Vim-F | TGCACGATGAAGAGATCCAGG |
| Mmu-Vim-R | CTTTCATACTGCTGGCGCAC |
| Mmu-Vinculin-F | GGTCTAGCAAGGGCAATGAC |
| Mmu-Vinculin-R | TGAATAAGTGCCCGCTTGGT |
