## Supplementary material for "The Epigenetic Regulator Histone Demethylase KDM5A is Activated and Pathogenic in a Mouse Model of Heart Failure": Table 1

**Table1:** **Echocardiographic parameters in WT, *Myh6-Cre*, *Kdm5a*^F/F^, and *Myh6-Cre:Kdm5a*^F/F^ mice at 2 months of age.**

| Groups | WT  (n=13) | *Myh6-Cre*  (n=15) | *Kdm5a*^F/F^  (n=12) | *Myh6-Cre* :*Kdm5a*^F/F^ (n=18) | P value |
| --- | --- | --- | --- | --- | --- |
| Sex (M/F) | 9/4 | 7/8 | 7/5 | 9/9 | Fisher exact test, P = 0.63 |
| Age (days) | 63.62±2.06 | 63.40±6.62 | 64±5.34 | 66.56±5.37 | Kruskal-Wallis,  P = 0.45 |
| BW (g) | 24.38±3.18 | 22.51±3.47 | 23.67±3.66 | 24.18±3.55 | Kruskal-Wallis,  P = 0.49 |
| HR (bpm) | 552±40 | 537±30 | 552 ±45 | 548±38 | One-way ANOVA,  P = 0.71 |
| IVST (mm) | 0.41±0.03 | 0.42±0.05 | 0.43±0.05 | 0.43±0.02 | Kruskal-Wallis,  P = 0.13 |
| PWT (mm) | 0.64±0.06 | 0.59±0.07 | 0.62±0.10 | 0.65±0.08 | One-way ANOVA,  P = 0.13 |
| LVEDD (mm) | 3.50± 0.27 | 3.33±0.30 | 3.36±0.40 | 3.43±0.26 | One-way ANOVA,  P = 0.43 |
| LVEDDi (mm/g) | 0.14±0.02 | 0.15±0.02 | 0.14±0.02 | 0.14±0.02 | Kruskal-Wallis,  P = 0.66 |
| LVESD (mm) | 2.17±0.32 | 2.18±0.25 | 2.13±0.31 | 2.21±0.30 | Kruskal-Wallis,  P = 0.91 |
| FS (%) | 37.66±4.38 | 34.85±3.80 | 36.50±4.11 | 35.66±5.45 | Kruskal-Wallis  P = 0.14 |
| EF (%) | 68.52±5.98 | 65.14±5.05 | 67.26±5.49 | 65.89±7.20 | Kruskal-Wallis,  P = 0.13 |
| LVM (mg) | 55.05±11.71 | 47.87±12.35 | 51.43±14.30 | 54.86±10.36 | One-way ANOVA,  P = 0.32 |
| LVMi(mg/g) | 1.82±0.37 | 1.73±0.33 | 1.78±0.44 | 1.92±0.32 | One-way ANOVA,  P = 0.46 |

Values are presented as mean ± SD. Normality was assessed using the Shapiro–Wilk test. Variables meeting normality assumptions were analyzed by one-way ANOVA with Tukey’s post hoc test, whereas non-normally distributed variables were analyzed using the Kruskal–Wallis test with Dunn’s correction. P values for the primary group comparison are shown. None of the parameters showed statistically significant differences; therefore, pairwise comparison P values were not significant. Abbreviations: M/F, male/female; HR, heart rate; BW, body weight; bpm, beats per minute; IVST, interventricular septal thickness; PWT, posterior wall thickness; LVEDD, left ventricular end-diastolic diameter; LVEDDi, LVEDD indexed to body weight; LVESD, left ventricular end-systolic diameter; FS, fractional shortening.
