## Supplementary material for "The Epigenetic Regulator Histone Demethylase KDM5A is Activated and Pathogenic in a Mouse Model of Heart Failure": Table 2

| Groups | WT  (n=14) | Myh6-Cre :Lmna^F/F^  (n=14) | *Myh6-Cre :Lmna^F/F^*  *:Kdm5a^F/F^*  (n=14) | P value |
| --- | --- | --- | --- | --- |
| Sex (M/F) | 8/6 | 7/7 | 10/4 | Fisher exact test  P = 0.62 |
| Age (days) | 20.00 ± 0.65 | 20.40 ± 0.91 ^a,c^ | 21.36 ± 0.50^a,b^ | One-way ANOVA,  P = 0.001 |
| BW (g) | 9.69 ± 2.20 | 7.86 ± 0.71^a^ | 9.1 ±1.07 | One-way ANOVA,  P = 0.007 |
| HR (bpm) | 516 ± 20 | 510 ± 15 | 549 ± 47^a,b^ | One-way ANOVA,  P = 0.003 |
| IVST (mm) | 0.39 ± 0.08 | 0.38 ± 0.08 | 0.40 ± 0.09 | One-way ANOVA,  P = 0.80 |
| PWT (mm) | 0.44 ± 0.03 | 0.41 ± 0.07 | 0.43± 0.08 | One-way ANOVA,  P = 0.47 |
| LVEDD (mm) | 2.70 ± 0.27 | 3.32 ± 0.30^a,c^ | 2.86 ± 0.36^b^ | One-way ANOVA,  P < 0.0001 |
| LVEDDi (mm/g) | 0.29 ± 0.06 | 0.44 ± 0.06^a,c^ | 0.32 ± 0.03^b^ | One-way ANOVA,  P < 0.0001 |
| LVESD (mm) | 1.51 ± 0.20 | 3.01 ± 0.35^a,c^ | 2.10 ± 0.42^a,b^ | One-way ANOVA,  P < 0.0001 |
| FS (%) | 43.96 ± 5.60 | 9.32 ± 4.60^a,c^ | 28.63 ± 7.38^a,b^ | One-way ANOVA,  P < 0.0001 |
| EF (%) | 81.92 ± 5.11 | 24.93 ± 10.40^a,c^ | 60.04 ± 12.47^a,b^ | One-way ANOVA,  P < 0.0001 |
| LVM (mg) | 20.40 ± 3.90 | 27.90 ± 7.80 ^a^ | 22.73 ± 5.88 | One-way ANOVA,  P = 0.007 |
| LVMi (mg/g) | 2.16 ± 0.41 | 3.53 ± 0.84^a,c^ | 2.50 ± 0.61^b^ | One-way ANOVA,  P < 0.0001 |

**Table 2: Echocardiographic parameters in WT, *Myh6-Cre* :*Lmna*^F/F^ and *Myh6-Cre* :*Lmna*^F/F^:*Kdm5a*^F/F^ at 3 weeks of age.**

Values are presented as mean ± SD. Normality was assessed using the Shapiro–Wilk test. Normally distributed variables were analyzed by One-way ANOVA followed by Tukey’s multiple-comparison test, whereas non-normally distributed variables were analyzed using the Kruskal–Wallis test with Dunn’s correction. Superscripts denote significant pairwise comparisons. **^a^** significant p value compared to WT, **^b^** Significant p value compared to *Myh6-Cre* :*Lmna*^F/F^, **^c^** Significant p value compared to *Myh6-Cre* :*Lmna*^F/F^:*Kdm5a*^F/F^. Abbreviations used: M/F - male/female, HR - heart rate, BW -body weight, bpm - beats per minute; IVST - interventricular septal thickness, PWT - posterior wall thickness, LVEDD - left ventricular end diastolic diameter, LVEDDi - LVEDD indexed to body weight, LVESD - left ventricular end systolic diameter, FS - fractional shortening.
