## Supplementary material for "The Epigenetic Regulator Histone Demethylase KDM5A is Activated and Pathogenic in a Mouse Model of Heart Failure": uncropped WBs

### Slide 1
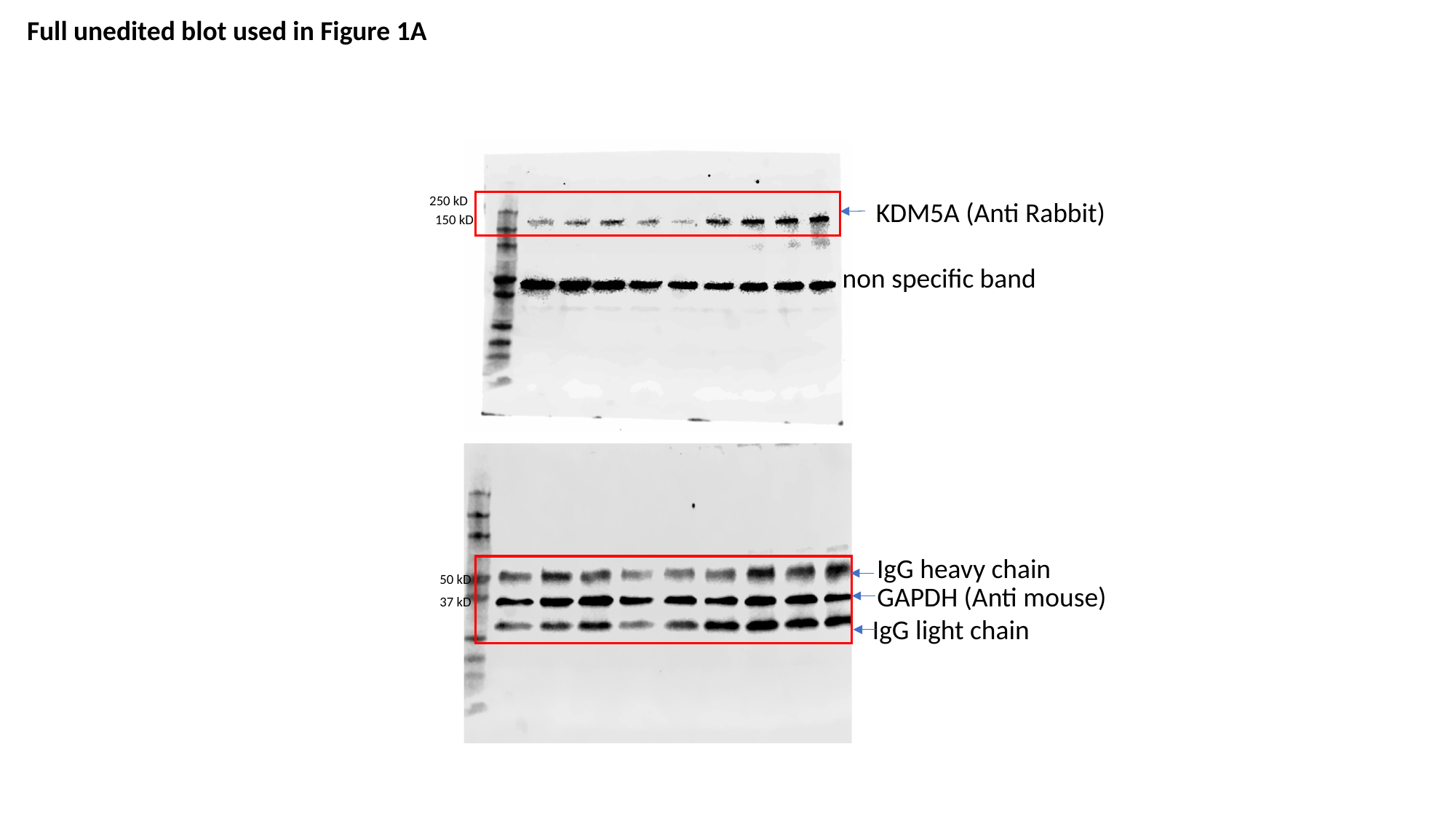

Full unedited blot used in Figure 1A
250 kD
KDM5A (Anti Rabbit)
150 kD
non specific band
IgG heavy chain
50 kD
GAPDH (Anti mouse)
37 kD
IgG light chain

### Slide 2
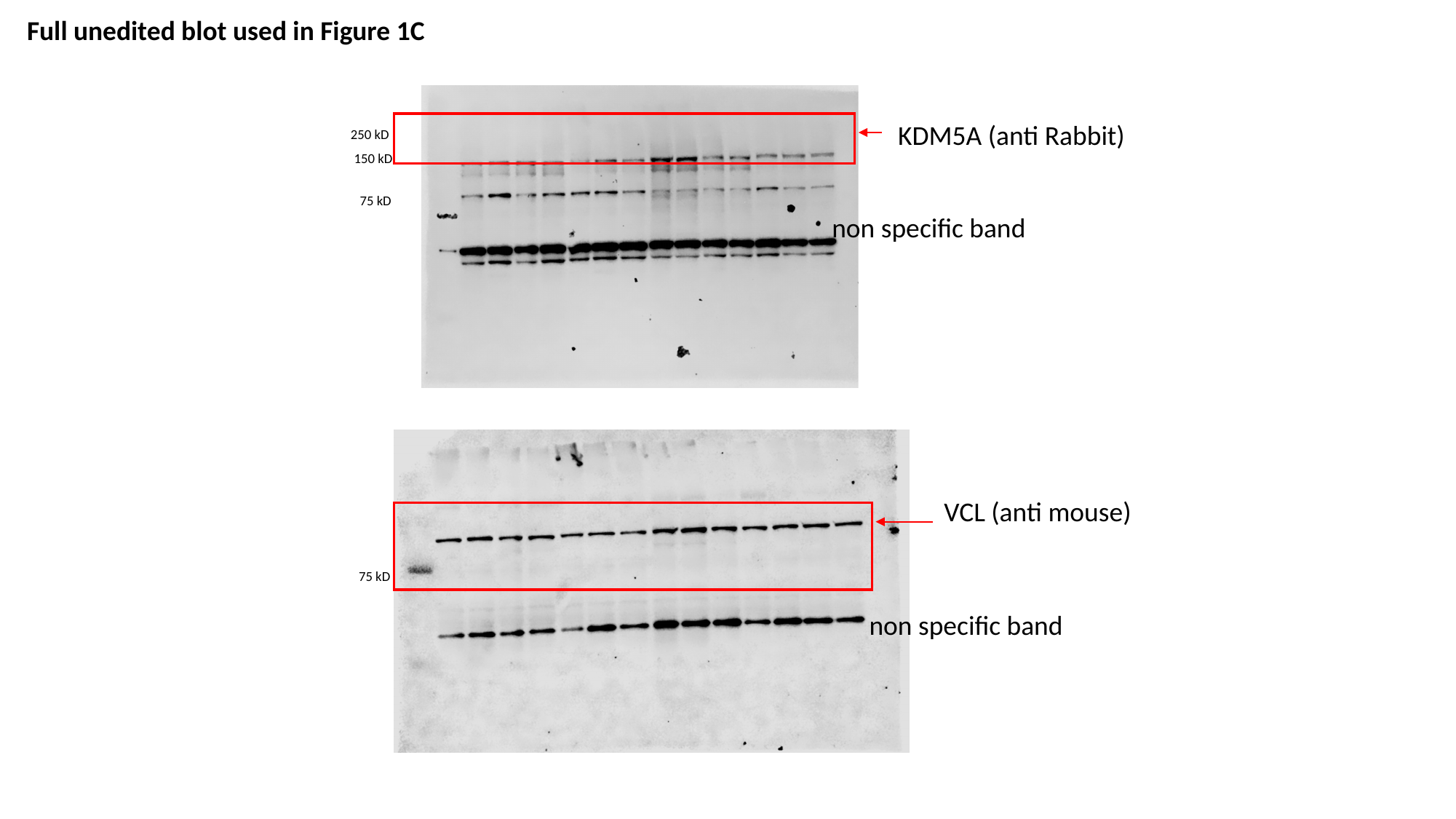

Full unedited blot used in Figure 1C
KDM5A (anti Rabbit)
250 kD
150 kD
75 kD
non specific band
VCL (anti mouse)
75 kD
non specific band

### Slide 3
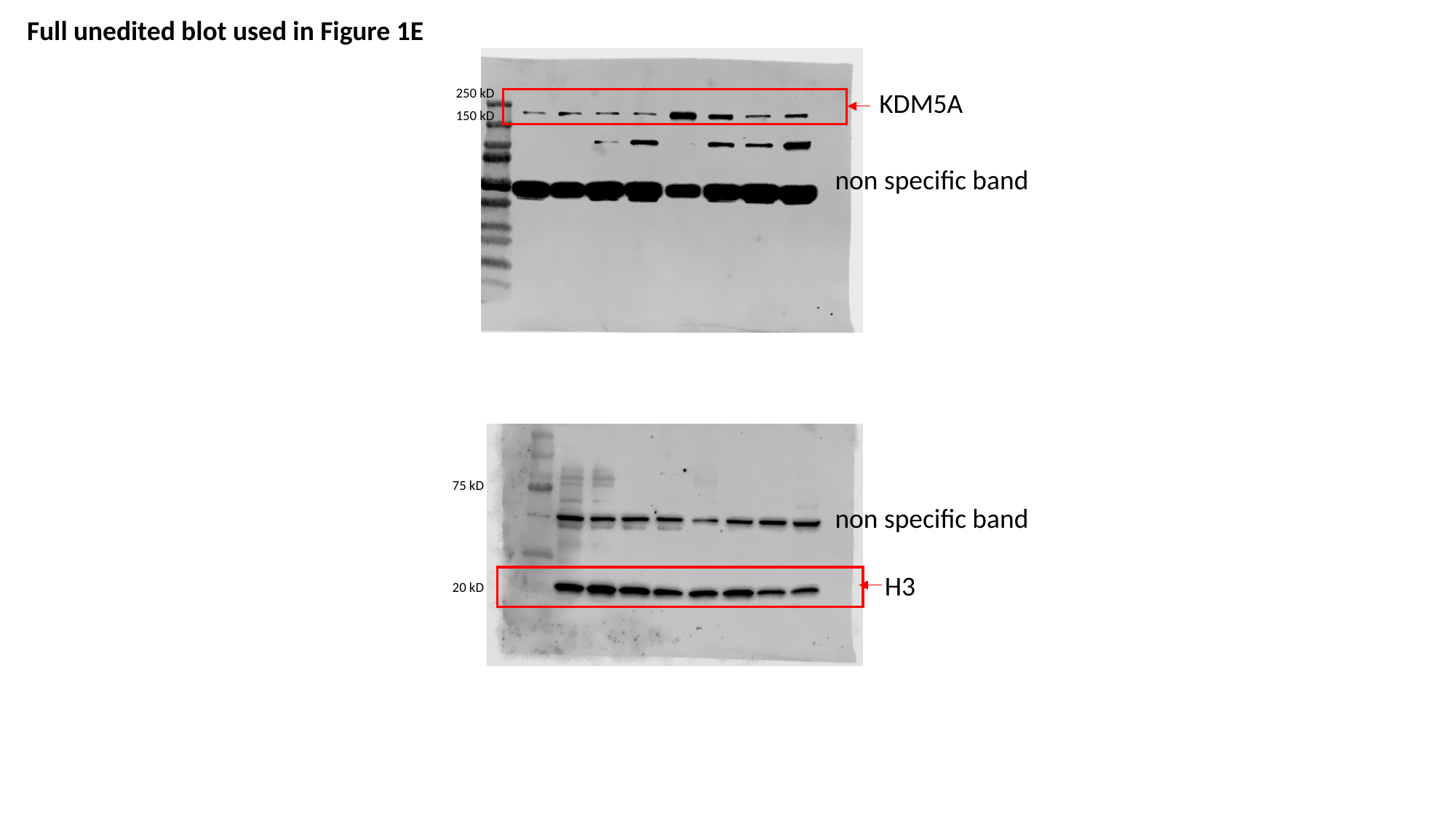

Full unedited blot used in Figure 1E
250 kD
KDM5A
150 kD
non specific band
75 kD
non specific band
H3
20 kD

### Slide 4
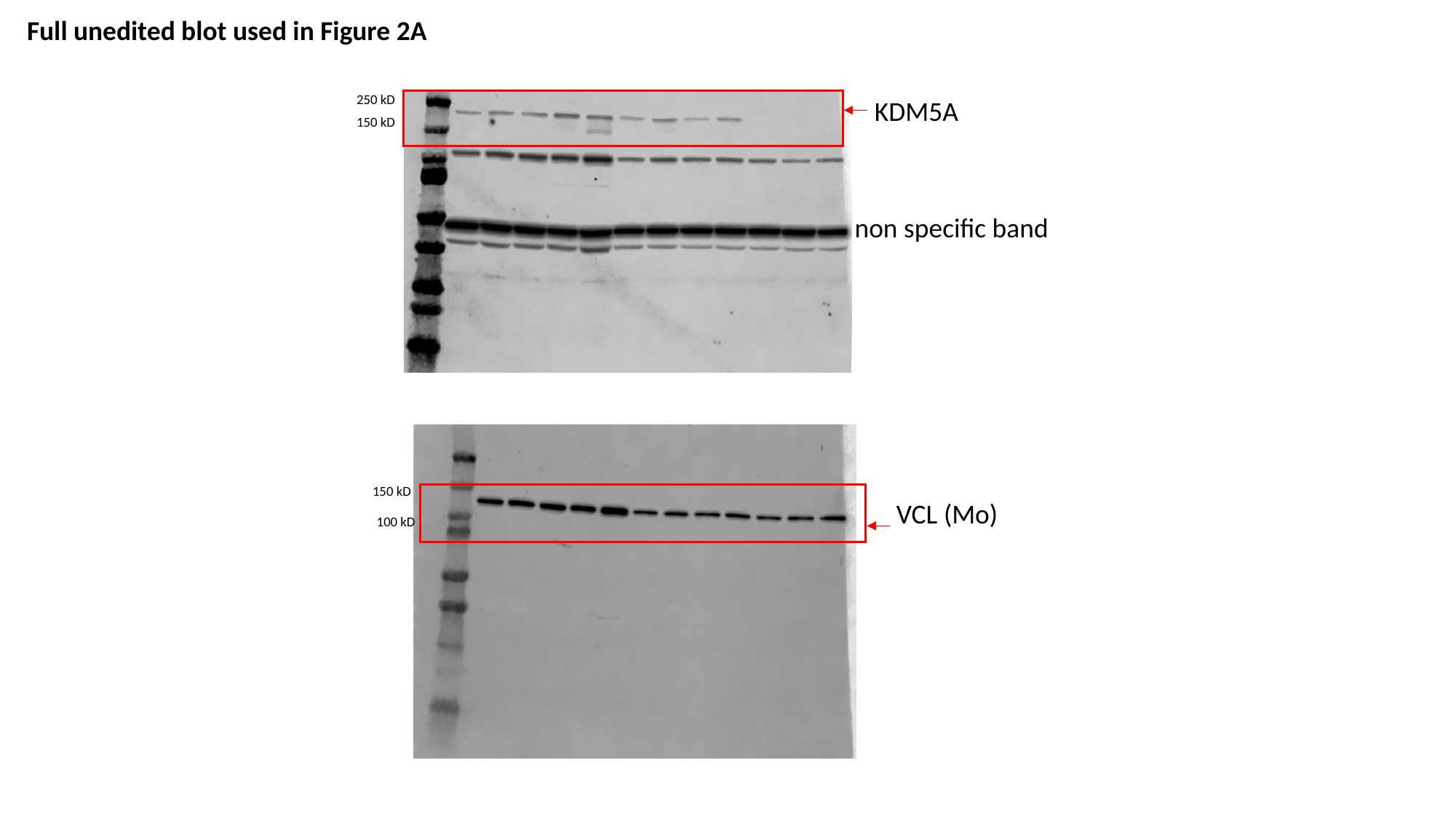

Full unedited blot used in Figure 2A
250 kD
KDM5A
150 kD
non specific band
150 kD
VCL (Mo)
100 kD

### Slide 5
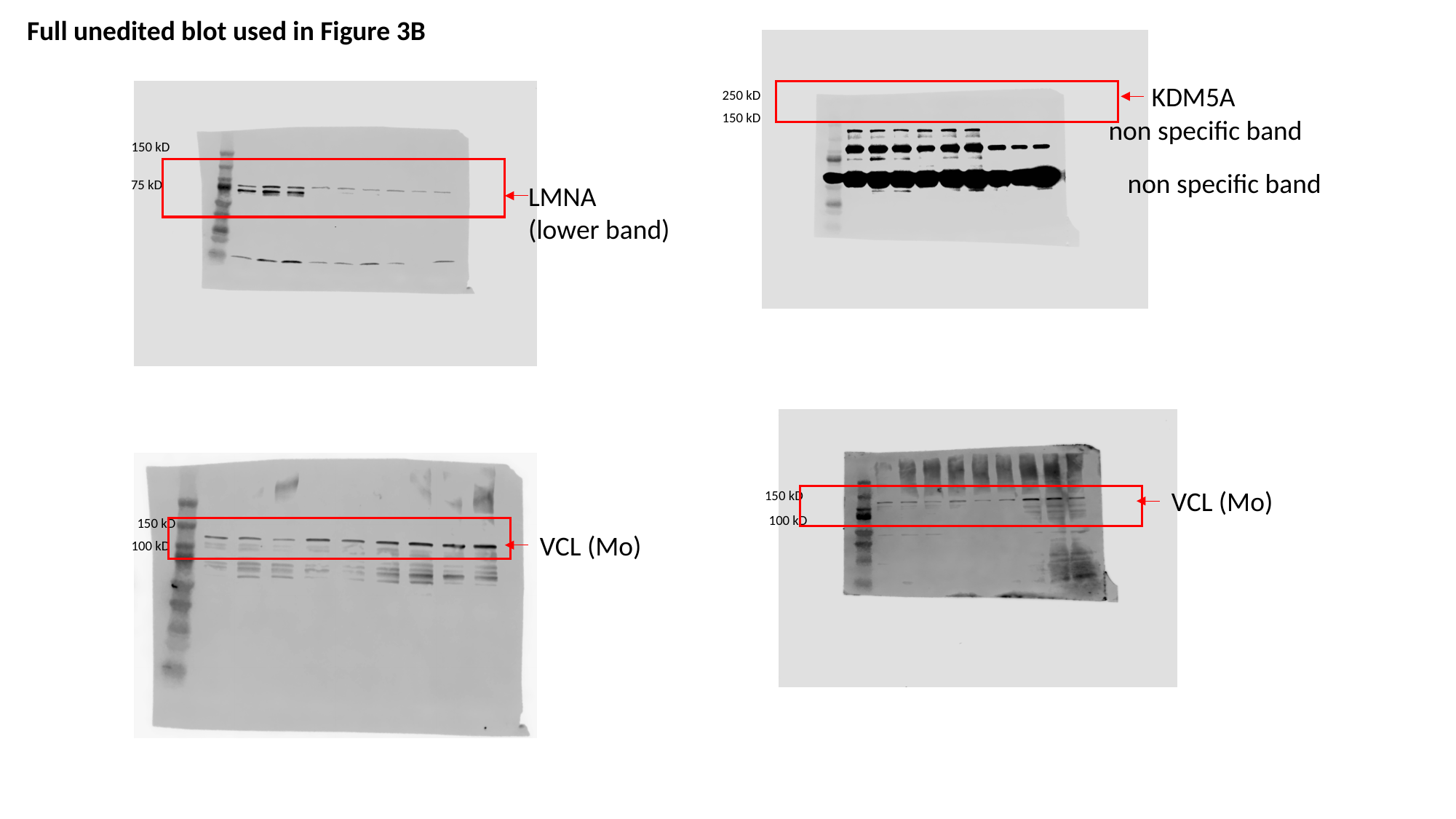

Full unedited blot used in Figure 3B
KDM5A
250 kD
150 kD
non specific band
150 kD
non specific band
75 kD
LMNA
(lower band)
VCL (Mo)
150 kD
100 kD
150 kD
VCL (Mo)
100 kD

### Slide 6
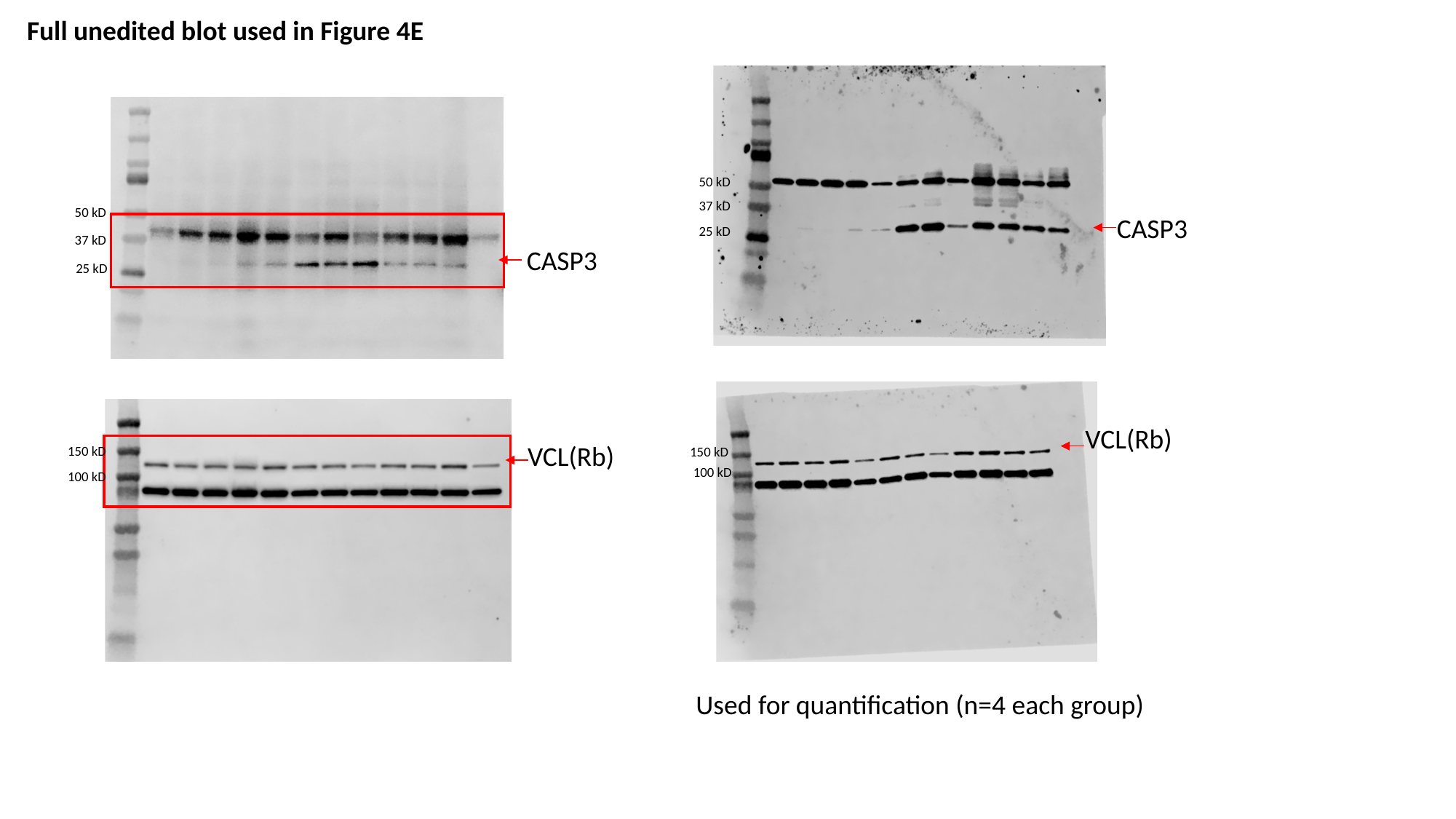

Full unedited blot used in Figure 4E
50 kD
37 kD
50 kD
CASP3
25 kD
37 kD
CASP3
25 kD
VCL(Rb)
VCL(Rb)
150 kD
150 kD
100 kD
100 kD
Used for quantification (n=4 each group)

### Slide 7
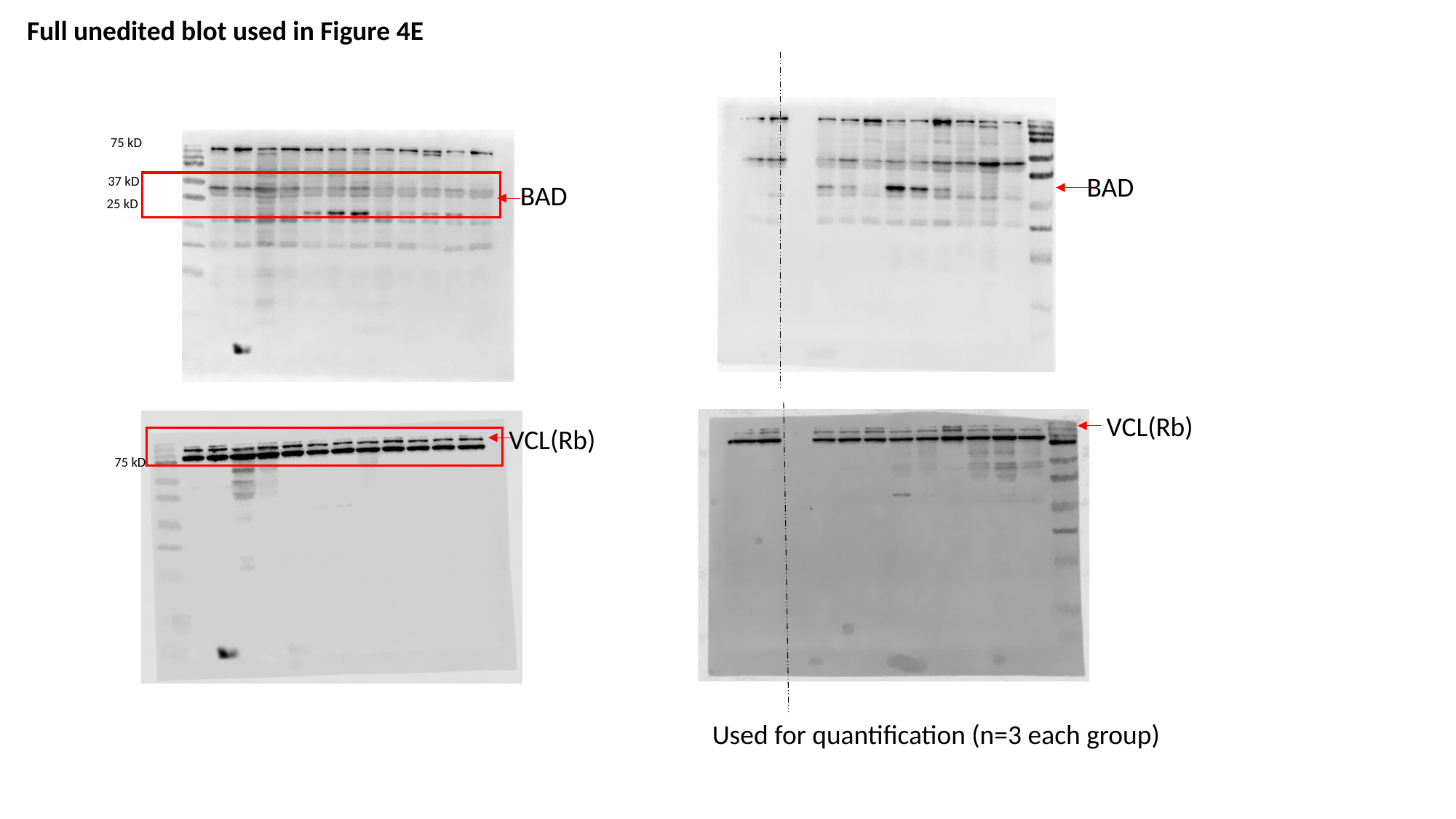

Full unedited blot used in Figure 4E
75 kD
BAD
37 kD
BAD
25 kD
VCL(Rb)
VCL(Rb)
75 kD
Used for quantification (n=3 each group)

### Slide 8
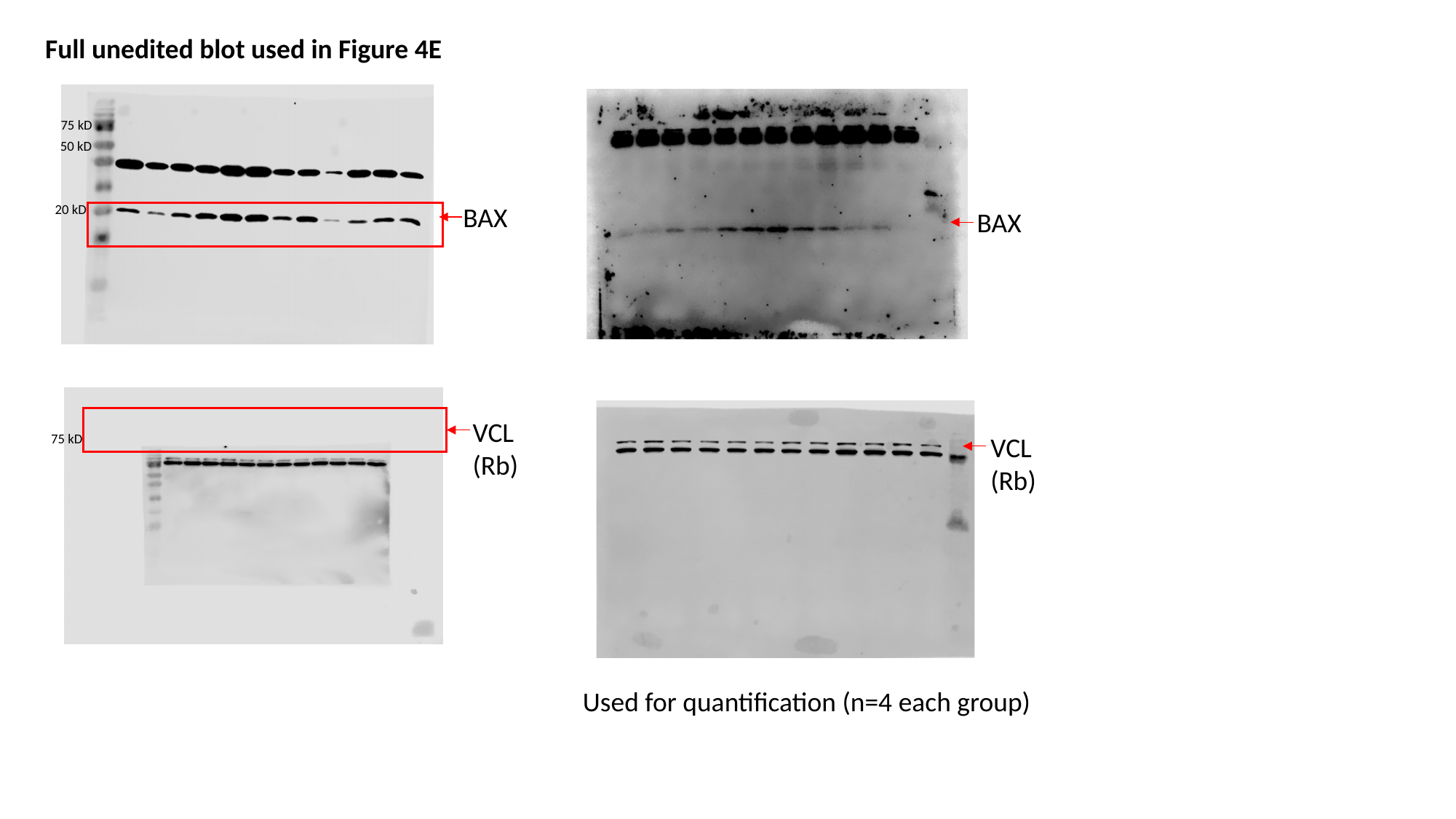

Full unedited blot used in Figure 4E
75 kD
50 kD
BAX
20 kD
BAX
VCL (Rb)
75 kD
VCL (Rb)
Used for quantification (n=4 each group)
